# Site-specific epigenetic marks in *Trypanosoma brucei* transcription termination, antigenic variation, and proliferation

**DOI:** 10.1101/2021.05.13.444086

**Authors:** Hee-Sook Kim

**Affiliations:** Public Health Research Institute, New Jersey Medical School, Rutgers, The State University of New Jersey, Newark, NJ 07103, USA

## Abstract

In *Trypanosoma brucei*, genes assemble into polycistronic transcription units (PTUs). Transcription termination sites (TTSs) hold deposition sites for three non-essential chromatin factors, histone variants (H3v and H4v) and a DNA modification (base J, a hydroxyl-glucosyl dT). Here, I found that H4v is a major sign for transcription termination at TTSs and readthrough transcription machineries progress until they encounter the next bidirectional transcription start site. While having a secondary function at TTSs, H3v is important for monoallelic transcription of telomeric antigen genes. The simultaneous absence of both histone variants leads to proliferation and replication defects, which are exacerbated by the J deficiency, accompanied by accumulation of sub-G1 population. Base J likely contributes to DNA replication and cell-cycle control. I propose that the coordinated actions of H3v, H4v and J function in concert for cellular fate determination and provide compensatory mechanisms for each other in chromatin organization, transcription, and replication.

## Introduction

*Trypanosoma brucei* is a parasitic protist that causes African sleeping sickness in humans and related diseases in animals, predominantly in sub-Saharan Africa. Disease transmission to mammals occurs by infected tsetse fly bites. To adapt in two different hosts, *T. brucei* undergoes stages of life-cycle specific differentiation. In insect vector, trypanosomes proliferate as a ‘procyclic form (PF)’ in the midgut and migrate to the fly’s salivary gland and differentiate into a non-proliferative metacyclic form. The infected fly injects the metacyclic form *T. brucei* into the mammalian host’s bloodstream. After differentiating into a ‘bloodstream form (BF)’, trypanosomes proliferate in the host’s bloodstream and extracellular spaces. BF trypanosomes are transferred during a blood meal and differentiate into a procyclic form and repeat the life cycle^1^. The surface of BF trypanosome is densely coated with a single type of Variant Surface Glycoprotein (VSG), which can trigger a strong antibody response in the mammalian host. But, trypanosomes can escape the host’s immune system with their reservoir of over 2,500 VSG genes by sequentially expressing one VSG gene at any given time, which drives disease persistence. Thus, transcriptional control of VSG genes and their genomic arrangement are key to *T. brucei* immune evasion mechanisms.

In *T. brucei*, groups of 10-100 genes are assembled in hundreds of Polycistronic Transcription Units (PTUs). Gene expression in *T. brucei* is controlled in three ways: 1) control of transcription initiation and termination at boundaries of PTUs, 2) processing of precursor RNAs (*trans*-splicing and polyadenylation) that generates mature mRNAs, and 3) regulating individual mRNA using regulatory information present in 3’
s UTRs and *trans*-acting elements like RNA-binding proteins^2–4^. PTU boundaries are demarcated by specific epigenetic marks in *T. brucei*. Transcription Start Sites (TSSs) feature acetylated H4 (H4K10ac), methylated H3 (H3K4me3), two of essential histone variants (H2Az and H2Bv), and base J (ß-D-glucosyl-hydroxymethyluracil, glucosyl-hmU)^5–7^, a *kinetoplastid*-specific DNA modification. Acetylated H2Az is required for H2Az deposition at TSSs. Depletion of HAT1, an acetyltransferase for H4, reduces total mRNA levels by 50%^8^. Deposition of H2Az at TSSs requires GT-rich sequence element present in TSSs^9^.

Transcription termination regulation in *T. brucei* remains less known. While TSSs are associated with essential histones, histone variants, and histone modifying enzymes, Transcription Termination Sites (TTSs) feature three non-essential factors, two of non-essential histone variants (H3v and H4v) and base J^5,7^. Base J modification uses a two-step process. J-Binding Protein-1 and 2 (JBP1, JBP2) mediate thymidine hydroxylation to generate a hydroxymethyl-dU (hmU) intermediate. J-associated Glucosyl Transferase (JGT) glucosylates hmU to generate base J^10,11^. Deletion of both JBP1 and JBP2 (JΔ) abolished base J modification in *T. brucei* but enable normal growth^10^, unlike in *Leishmania* species with essential JBP1 functionsl^12^. Thus, base J may function differently than in the closely related *kinetoplastid* parasites. Approximately 50% of base J localizes at PTU boundaries of PTUs in *T. brucei*, but, only 1% of base J is chromosome internally primarily at TTSs in *Leishmania*^13–16^. J depletion significantly increased transcription readthrough at TTSs in *Leishmania*^17,18^. However, J null trypanosomes showed minor transcription termination defects with increased antisense transcripts concentrated near TTSs^19,20^. In both organisms, no significant transcription termination defects were observed in the H3v KO single mutant^19,21^. However, antisense transcript levels were higher in trypanosome cells lacking both H3v and J than J null cells^19,20^, indicating that H3v also contributes to transcription termination in *T. brucei*.

The last PTU in some megabase chromosomes houses a VSG, a trypanosome surface antigen gene. The relationship between H3v and J is mirrored in transcription regulation at these telomeric PTUs^19^. Allelic exclusion of VSG among the 2,500 VSG genes and periodic switching of the expressed allele allows trypanosomes to evade the host immune response, a phenomenon known as the antigenic variation. VSG genes occur at four types of chromosomal loci^22–27^: 1) Bloodstream-form Expression Site (BES), 2) Metacyclic Expression Site (MES), 3) Minichromosome (MC), and 4) subtelomeric region. *T. brucei* genome has about 15-20 BESs. A BES, a special telomeric PTU, contains a promoter associated with RNA pol I transcription (not pol II), several of Expression-Site Associated Genes (ESAGs), 70-bp repeats, and a VSG gene located immediately upstream of telomere repeat. There are 6 MESs, each containing a VSG immediately upstream of telomere repeat. MES VSGs are expressed specifically in the metacyclic stage by RNA pol I. *T. brucei* has about 60 small linear chromosomes called minichromosomes (about 30-150kb in size). MCs are organized with an inverted 177-bp repeat in the middle and telomeres at the ends. Some MCs have a VSG located immediately upstream of telomere repeat, but no promoter is present^24^. The remaining VSGs are found in arrays without promoters at subtelomeric regions. Only one BES promoter is transcriptionally active, while the others are repressed to enable monoallelic expression of VSG. A new VSG can be activated by changes in transcriptional status or genetic rearrangement that moves a silent VSG to the active VSG location (gene conversion)^28^. Trypanosomes coated with a new VSG are undetected by host antibody defenses tuned to prior VSG. H3v and base J reside in telomere repeats in *T. brucei*^5,7^, but only H3vΔ showed increased derepression of telomeric silent VSGs^19^. Level of silent VSG derepression was higher in H3vΔ JΔ double mutant compared to H3vΔ^19^, indicating that VSG silencing requires H3v with minor contribution by J.

In contrast to TSS chromatin marks, TTS marks are non-essential^5,10,19,29^. I hypothesized that H3v, J and H4v will perform redundant functions at TTSs. If this supposition were true, deleting all three marks will induce severe termination defects and result in cell lethality or sickness. Here, our study indicates that H3v, H4v and base J did show synthetic lethal genetic interaction due to uncontrolled cell division with stalled replication, not transcription termination defects. H4v is a major mark of transcription termination but H4vΔ cells grow normally. While having a secondary function at TTSs, H3v is important for monoallelic transcription of VSG genes. The absence of both histone variants promoted cell growth and replication defects, which increased in the absence of base J. Thus, the coordinated actions of H4v, H3v and J in chromatin determine cell fate.

## Results

### Synthetic lethal genetic interaction among TTS-associated chromatin marks

I predicted that successful transcription termination in *T. brucei* requires three non-essential chromatin marks (H3v, H4v, and base J) that coincide at TTSs, which may functionally “signal” RNA polymerase II to stop. All experiments were performed in BF trypanosomes, as J modification is detected only in BF stage. I generated individual single gene knockout (KO) mutants, H3vΔ^19^, H4vΔ, JΔ^19^, using an established Cre-loxP system^30^. Although JΔ was generated by sequentially deleting both alleles of JBP1 and JBP2 genes, this study will refer to them as a single mutant hereafter, as J is one of three TTS marks. Briefly, all parental cell lines contain a Cre recombinase gene whose expression can be induced by adding tetracycline (‘Tet-on’). Gene deletion cassettes have upstream and downstream homology sequences and a floxed selection marker in the middle. Expression of Cre by Tet addition can remove floxed selection markers, so that they can be reused. JΔ strain without markers serve to generate H3vΔ JΔ^19^ and H4vΔ JΔ strains. Both double mutants and the H4vΔ mutant grew normally and showed no morphological abnormalities. To generate a H4vΔ H3vΔ mutant, I transfected a H4v KO vector in the H4vΔ/+ heterozygous H3vΔ strain. The six viable clones grew slower than WT. Four clones showed an unusual growth pattern, as cell growth temporarily arrested before growth resumed (Supplementary Fig. 1, arrows). These double KO cells may have experienced replication challenges but recovered through epigenetic adaptations. To avoid any obscurity cause by unknown events occurring during prolonged culturing and analyze the mutant phenotypes immediately after the knockout, I generated a conditional KO mutant, a H4vΔ strain with a floxed H3v-Ty1, termed conditional double KO (DKO). Cre expressed by tetracycline can remove the floxed H3v-Ty1 allele to produce H3vΔ H4vΔ cells (DKO + Tet) (Fig. 1A). No viable clones emerged after deleting the remaining H4v allele in the H4vΔ/+ heterozygous H3vΔ JΔ strain. So, I generated a conditional triple KO mutant, a H4vΔ JΔ strain with a floxed H3v-Ty1, termed conditional triple KO (TKO). Genotypically, TKO is a quadruple null of H3v, H4v, JBP1 and JBP2 genes.

**Figure 1.**
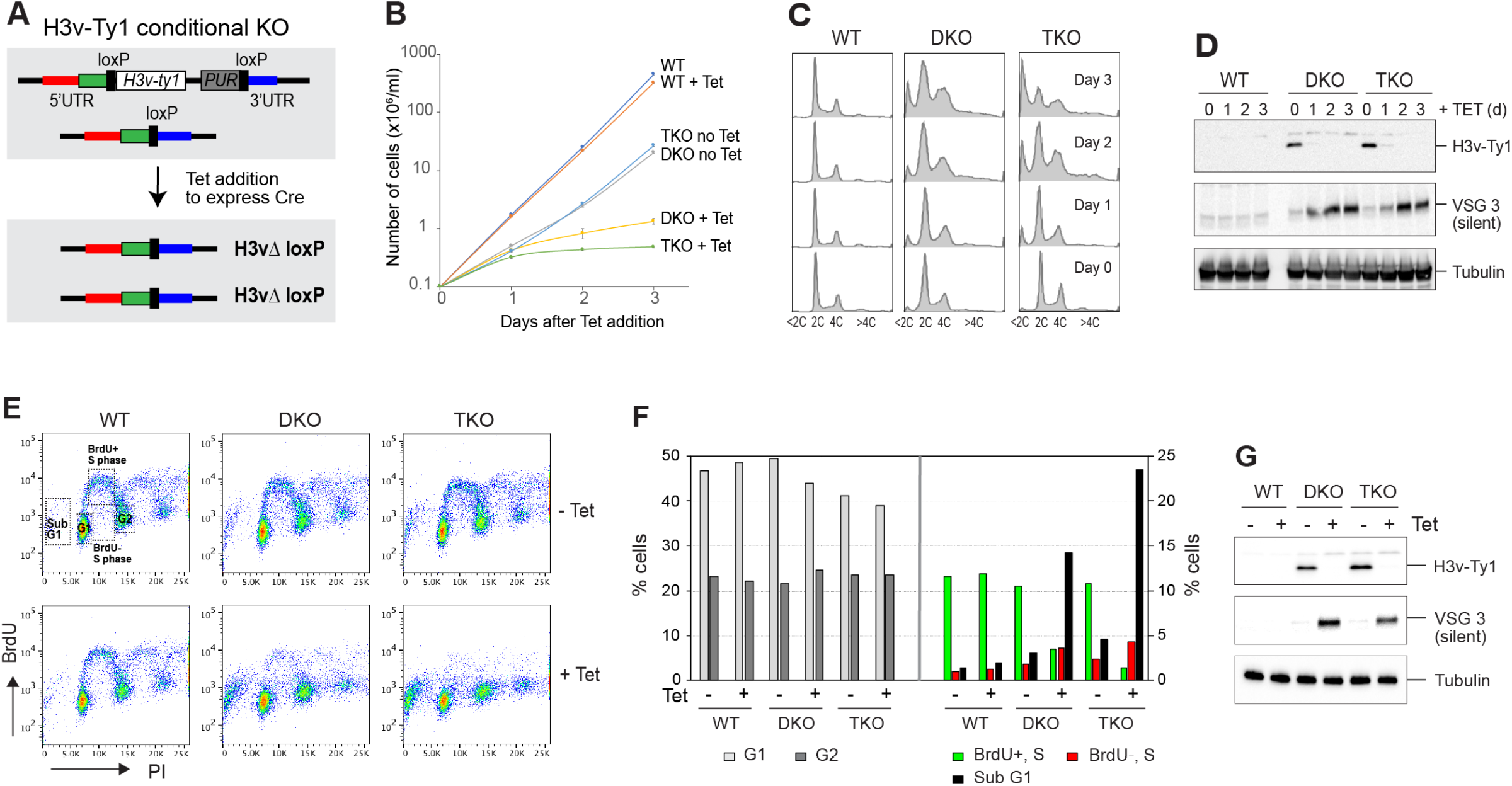
Genetic interaction between H3v, H4v and base J in proliferation, cell-cycle progression and DNA replication. **(A)** Strategy to generate a conditional double KO (DKO, floxed H3v-Ty1 in H4vΔ strain) and a conditional triple KO (TKO, floxed H3v-Ty1 in H4vΔ JΔ strain). The strain has one H3v allele deleted and the other H3v allele replaced with a floxed H3v-Ty1 and PUR marker. Cre protein expressed by tetracycline removes the floxed H3v-Ty1-PUR in H4vΔ or H4vΔ JΔ strain, generating H3vΔ H4vΔ (DKO + Tet) or H3vΔ H4vΔ JΔ (TKO + Tet) cells. **(B)** Cell growth in DKO and TKO mutants upon Tet induction. WT, DKO and TKO strains were treated with tetracycline and cell count was monitored daily in triplicate. **(C)** Cell-cycle progression in DKO and TKO mutants upon Tet induction. Cells from (B) were fixed, stained with Propidium Iodide (PI) and analyzed by flow cytometry. **(D)** Depletion of H3v-Ty1 protein and derepression of silent VSG3 protein in DKO and TKO mutants upon Tet induction. Denatured whole cells from (B) were analyzed by western blot. Tubulin served as a loading control. **(E)** BrdU incorporation in DKO and TKO mutants upon Tet induction. WT, DKO and TKO strains untreated or treated with tetracycline for 2 days were pulse-labeled with 500 μM BrdU for 40 min and fixed. Cells were stained with PI (bulk DNA) and anti-BrdU-Alexa 488, and then analyzed by flow cytometry. **(F)** Quantification of BrdU-positive or negative S phase population and, G1, G2, and sub-G1 cell population. **(G)** Western blot control for experiment E.

After adding Tet, both DKO and TKO cells exhibited severe growth defects and abnormal cell-cycle distribution (greater in the Tet-treated TKO) (Fig. 1B & 1C). Cell populations containing less than 2C DNA content (sub-G1) significantly increased after removing H3v in the conditional TKO strain (Fig 1C). Levels of VSG3, a silent VSG protein, increased after Tet treatment in DKO and TKO (Fig. 1D). The DKO double mutant grew more poorly than those generated by transfection, suggesting that cells compensate with the loss of H3v and H4v marks in different mechanisms over time.

I observed increased sub-G1 population in *Tb*MCM-BP depleted cells^31,32^, similar to Tet-treated TKO. *Tb*MCM-BP depletion also caused a DNA replication defect^32^. To determine if replication problems triggered these cell-cycle defects in TKO cells, I measured the efficiency of DNA synthesis using bromo-2’deoxyuridine (BrdU) incorporation. Tet-untreated WT, DKO and TKO showed a normal pattern of BrdU labeling. BrdU incorporating S phase population was decreased in Tet-treated DKO and this reduction was more pronounced in Tet-treated TKO (Fig. 1E, F). I posit that sub-G1 population in Tet-treated TKO cells arise as mutant cells enter cytokinesis before completing DNA replication.

### H4v drives transcription termination with H3v-J contributions

JΔ and H3vΔ JΔ mutants showed mild transcription termination defects and wild type growth rates^19,20^. I speculated that these effects reflected the compensatory presence of H4v, such that simultaneous deletion of H3v, H4v, and J would cause stronger termination defects. So, I performed stranded RNA-seq experiment with triplicated cultures of WT and 9 KO mutants, including JΔ, H3vΔ, H4vΔ, H3vΔ JΔ, H4vΔ JΔ, DKO without or with Tet treatment, and TKO without or with Tet treatment. Strains were confirmed by PCR and/or western blot and flow cytometry (Supplementary Fig. 2 & 3). Stranded total RNA was prepared by rRNA removal to avoid the loss of antisense transcripts that are not polyadenylated. Sequence reads were analyzed with Bowtie 2^33^ and Seqmonk (Babraham Bioinformatics).

Stranded RNA-seq reads were aligned to the Lister 427 genome. 11 megabase chromosomes were analyzed with sliding windows (5 kb bin, 1kb step). Reads Per Million mapped reads (RPM) values were generated separately from forward and reverse reads and fold changes compared to WT were plotted. Chromosome 10 (Fig. 2) and 7 (Supplementary Fig. 5) are shown as examples. TSSs are shown with H4K10ac peaks^5^. Forward reads mapping to reverse PTUs (blue) (transcription moving reverse direction) represent sense transcription, and forward reads mapping to forward PTUs (red) (transcription moving forward) represent antisense transcription. Reverse reads mapping to forward PTUs (red) represent sense and to reverse PTUs represent antisense transcription. Transcript level was unchanged in JΔ and H3vΔ, compared to WT (Fig. 2A). H4vΔ (pink lines) showed an approximately 2-fold increase in antisense transcription throughout chromosome 10. In the region for PTU assembly in head-to-tail (HT) organization (Fig. 2), all PTUs have the same increase in antisense transcript levels. The data indicate that H4v is an important signal for transcription stop. RNA pol II machinery that missed its stop sign progresses until encountering a bidirectional transcription start site, where directionality of RNA pol II should be determined^9,34,35^.

**Figure 2.**
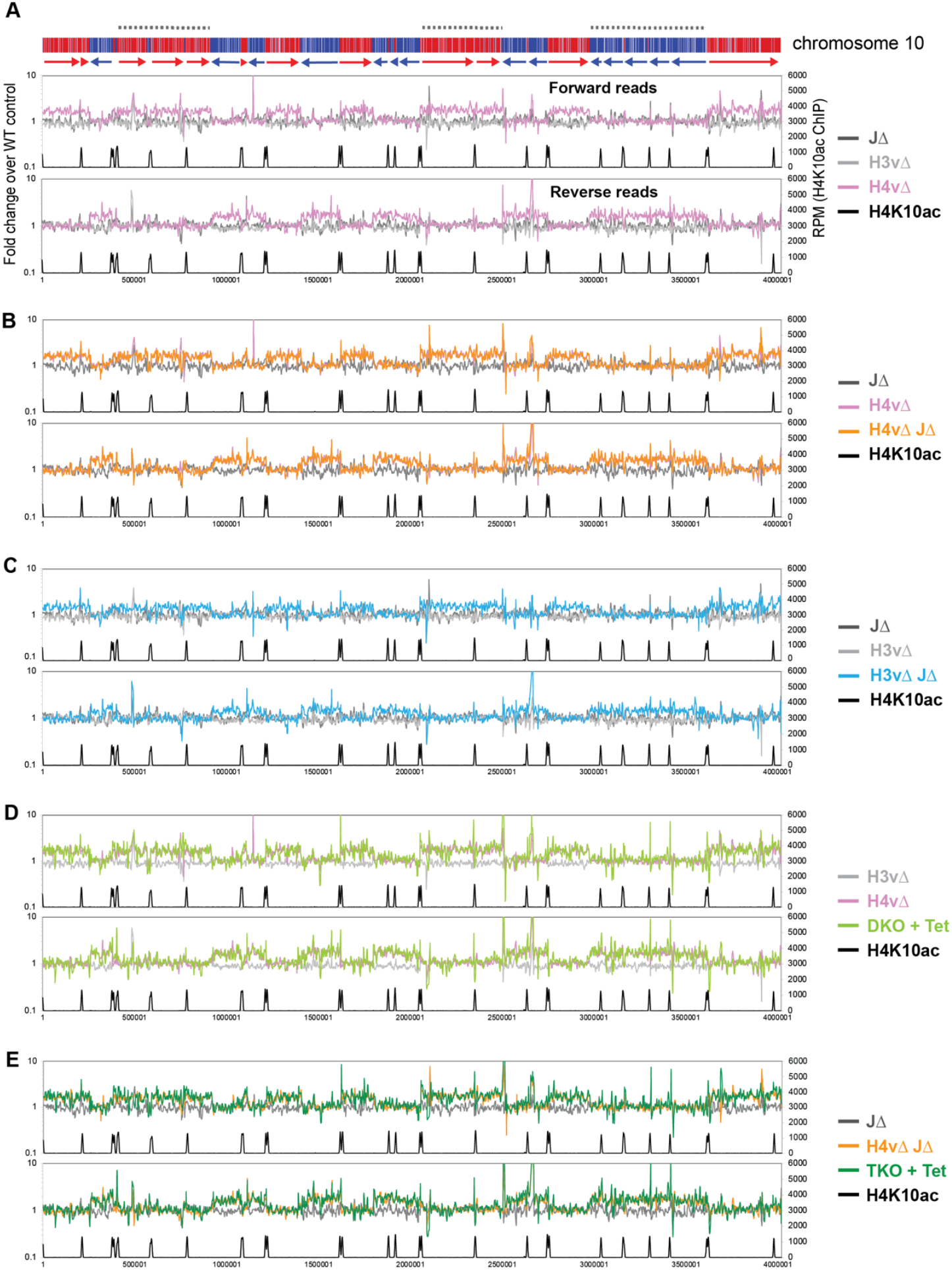
H4v is the major transcription termination signal with a secondary role of H3v-J. Global transcription was examined by rRNA-depleted stranded RNA-seq and mapping the sequence reads to the *T. brucei* Lister 427 genome. Forward and reverse reads were analyzed separately with sliding windows (5kb bin, 1kb step). Fold changes of RPM values between WT and each of KO mutant are plotted for all chromosomes. Chromosome 10 is shown as an example. Diagram of chromosome 10 is shown on top: PTUs are shown in red and blue bars with arrow heads indicating transcription direction. Dotted lines are regions containing multiple HT PTUs in the same direction. Primary y-axis is a fold change of RPM values of each mutant compared to WT. Secondary y-axis is RPM values obtained from H4K10ac ChIP seq. H4K10ac ChIP raw reads obtained from the Cross lab^5^ were mapped to the Lister 427 genome with the Bowtie 2. Following KO mutants are compared in each plot **(A)** JΔ, H4vΔ, and H3vΔ. **(B)** JΔ, H4vΔ, and H4vΔ JΔ. **(C)** JΔ, H3vΔ, and H3vΔ JΔ. **(D)** H3vΔ, H4vΔ, and Tet-treated DKO (H3vΔ H4vΔ). **(E)** JΔ, H4vΔ JΔ, Tet-treated TKO (H3vΔ H4vΔ JΔ).

No further increase in antisense transcription was found in H4vΔ JΔ compared to H4vΔ strain (Fig. 2B). This same pattern of antisense-transcription increase occurred in H3vΔ JΔ strain (Fig. 2C), but to a lesser degree. Two possible mechanisms could underlie this. First, H4v and H3v-J independently control transcription termination, which predicts that an antisense transcription increase would be higher in triple KO than in H4vΔ mutant (additive or synergistic). Second, H4v is the major signal for termination with assistance from H3v-J, so H4vΔ H3vΔ and H4vΔ H3vΔ JΔ should show similar levels of antisense transcription as the H4vΔ. As shown in Fig. 2D and 2E, the increase in antisense transcription was comparable between H3vΔ H4vΔ (DKO + Tet) and H4vΔ and between H3vΔ H4vΔ JΔ (TKO + Tet) and H4vΔ JΔ. The same pattern occurred along all chromosomes (Supplementary Fig. 5). Therefore, H4v is the major chromatin mark for transcription termination with secondary contribution from H3v-J.

RPKM values of all, sense, or antisense reads mapping to 8,428 CDSs (excluding those located in subtelomeric region), are presented in Supplementary Fig. 6A. Mean values of antisense transcription increased about 1.44 (H3vΔ JΔ), 1.64 (H4vΔ), 1.75 (DKO + Tet) and 1.77-fold (TKO + Tet) compared to WT, while sense transcription did not significantly change. Location of CDSs with upregulated antisense transcription confirmed that the antisense transcription increase is global and evenly distributed throughout all PTUs (Supplementary Fig. 6B).

### TTS chromatin marks affect polyadenylation of antisense transcripts

We demonstrated that antisense transcription was significantly increased at and near TTSs in JΔ in polyA selected stranded RNA-seq experiments^19^. In the rRNA deletion method here, antisense transcription showed minimal alterations in JΔ mutant cells (Fig. 2). This suggests that TTS chromatin marks may be involved also in polyadenylation of antisense RNA. So, I compared WT stranded RNA-seq data prepared by polyA selection and rRNA depletion. Reads mapped to the Lister 427 genome were analyzed with sliding windows (5 kb bin, 1kb step), as in Fig. 2. RPM values for each method were plotted for chromosome 10 (Fig. 3A). Forward reads mapping represents sense transcription for reverse PTUs (blue) and antisense for forward PTUs (red), and vice versa for reverse reads mapping. For sense transcription, I found no change between samples prepared by polyA selection and rRNA depletion, but a higher level of antisense transcripts (about 10-100-fold) in rRNA depleted samples. This result indicates that only a small fraction of antisense transcripts undergoes polyadenylation.

**Figure 3.**
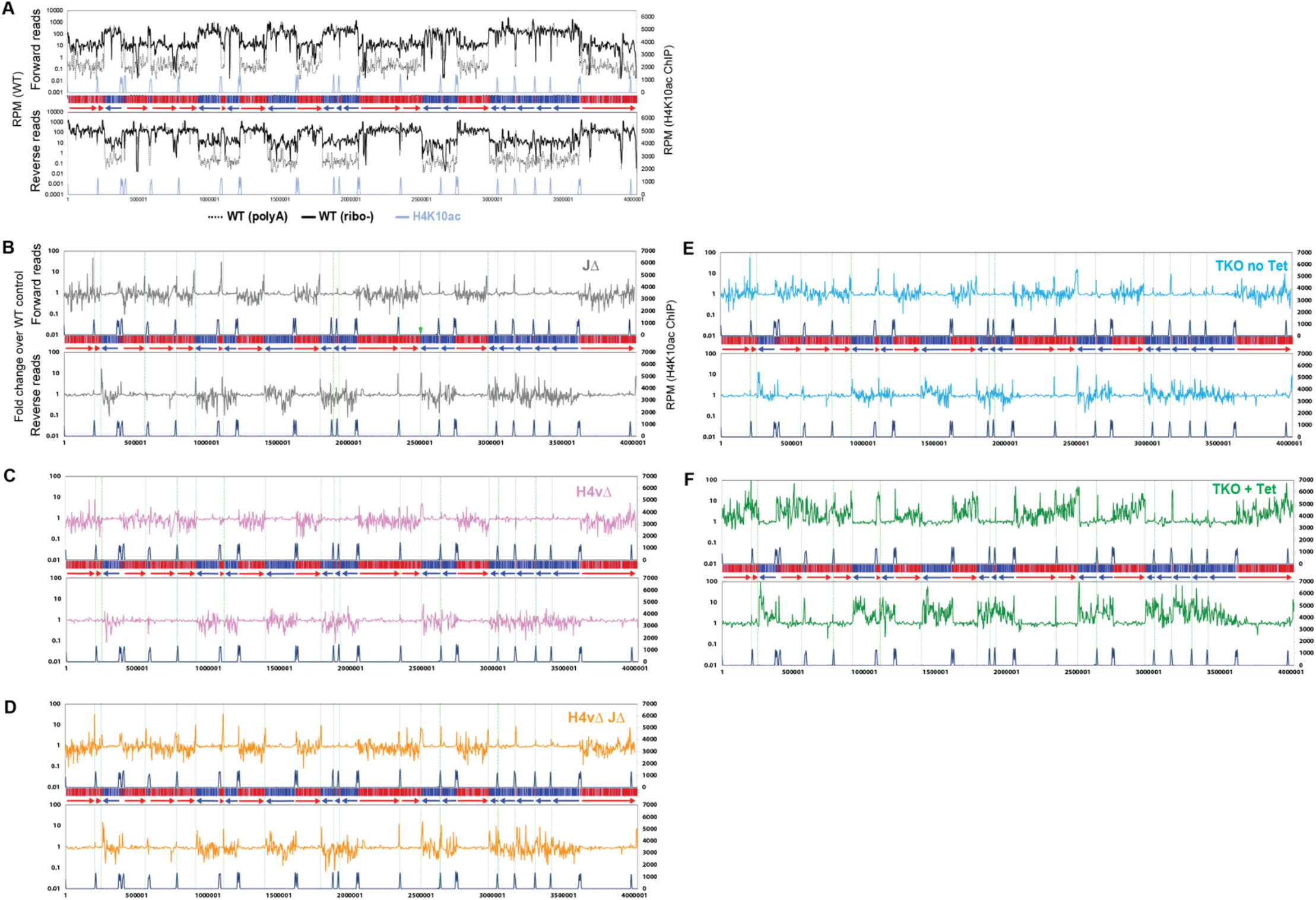
TTS-chromatin marks affect polyadenylation of antisense transcripts. Global transcription was examined by polyA-selected stranded RNA-seq and mapping to the Lister 427 genome. **(A)** Forward and reverse reads were analyzed separately with sliding windows (5kb bin, 1kb step). RPM values from WT RNA-seq prepared by polyA selection (dotted black line) were compared with those from WT rRNA-depleted RNA-seq (solid black line). RPM values were plotted for all chromosomes. Shown chromosome 10 as an example. Diagram of chromosome 10 is depicted in the middle of forward and reverse plots. **(B-F)** Fold changes between wild type and each KO mutant are plotted. Shown chromosome 10: JΔ (B), H4vΔ (C), H4vΔ JΔ (D), TKO without Tet (E), TKO with Tet (F). Green dotted lines indicate locations of TTSs.

The increase of polyadenylated antisense transcripts (spiking peaks near TTSs: green lines in Fig. 3B) was observed near TTSs in J Δ, consistent with prior results^19^. This was also observed in H4vΔ JΔ and TKO without Tet (Flag-H4v cKO in H3vΔ JΔ) (Fig. 3D, E). Interestingly, Tet-treated TKO cells undergo an increase in polyadenylated antisense transcripts that were not only localized at TTSs, but also at all PTU regions (Fig. 3F). H4vΔ alone did not significantly change the level of polyadenylated antisense transcripts (Fig. 3C). The data suggest that H4v is required for stopping the RNA pol II at TTSs, but all three TTS marks are required to stop the polyadenylation of non-coding RNA produced by readthrough transcription. Transcriptome analysis of 8,428 CDSs showed that antisense transcription was 1.69-fold higher in the triple KO mutant compared to WT (Supplementary Fig. 6C). Overall, I conclude that TTS chromatin marks control transcription termination and also define the junction of coding and non-coding regions within a long precursor RNA by preventing polyadenylation of antisense RNA. Statistical analyses for all transcriptome data are summarized in Supplementary Table 7.

### TTS-chromatin marks control VSG transcription in a synergistic manner

While chromosome internal TTSs have all three marks, telomeres are enriched with H3v and base J, but not H4v^5,7,29^. Transcription is repressed at telomeric regions by heterochromatin structure in *T. brucei* as in other eukaryotes^36–38^, and transcription of telomeric VSG utilizes RNA pol I^39^, not pol II. Therefore, I speculated that the genetic requirement for VSG transcription control would be different from that for RNA pol II-mediated transcription control. Four types of VSG locations are depicted in Fig. 4A. A single VSG is transcribed from one BES (telomeric PTUs with pol I promoter in 10-60kb length)^25,26,39,40^. Additional genomic locations that contain VSGs are also telomeric (MES^26,27^ or minichromosomes^24^) or subtelomeric, chromosome-internal VSGs (mostly partial genes or pseudogenes). Only one BES is transcriptionally active at a time, while the remaining BESs are silent. Sequence reads were mapped to the VSGnome^26^ and LOG2(RPKM) values of VSG CDSs were compared between WT and KO mutants (Fig. 4). H4vΔ or H4vΔ JΔ cells exhibited no significant changes in levels of silent BES and MES VSG transcripts (Fig. 4B, C). Mutants lacking H3v showed increased levels of silent BES and MES VSGs like DKO+Tet, TKO+Tet > H3vΔ JΔ > H3vΔ (high to low). Given that H4vΔ and H4vΔ JΔ showed no change in VSG silencing and H4v does not localize at the telomere, the synergistic upregulation of silent VSGs in Tet-treated DKO and TKO cells was puzzling. H4v could bind telomere but are not detectable, or H4v may bind telomeres only in the absence of H3v. Transcription at the entire BES units were also examined. Interestingly, only those genes adjacent to promoter or telomere (VSG) were significantly affected by H3v, H4v and J (Supplementary Fig. 7). Telomere unprotection in Tet-treated DKO or TKO cells may cause disruption of silent BES heterochromatin structure, leading to the loss of promoter and VSG silencing.

**Figure 4.**
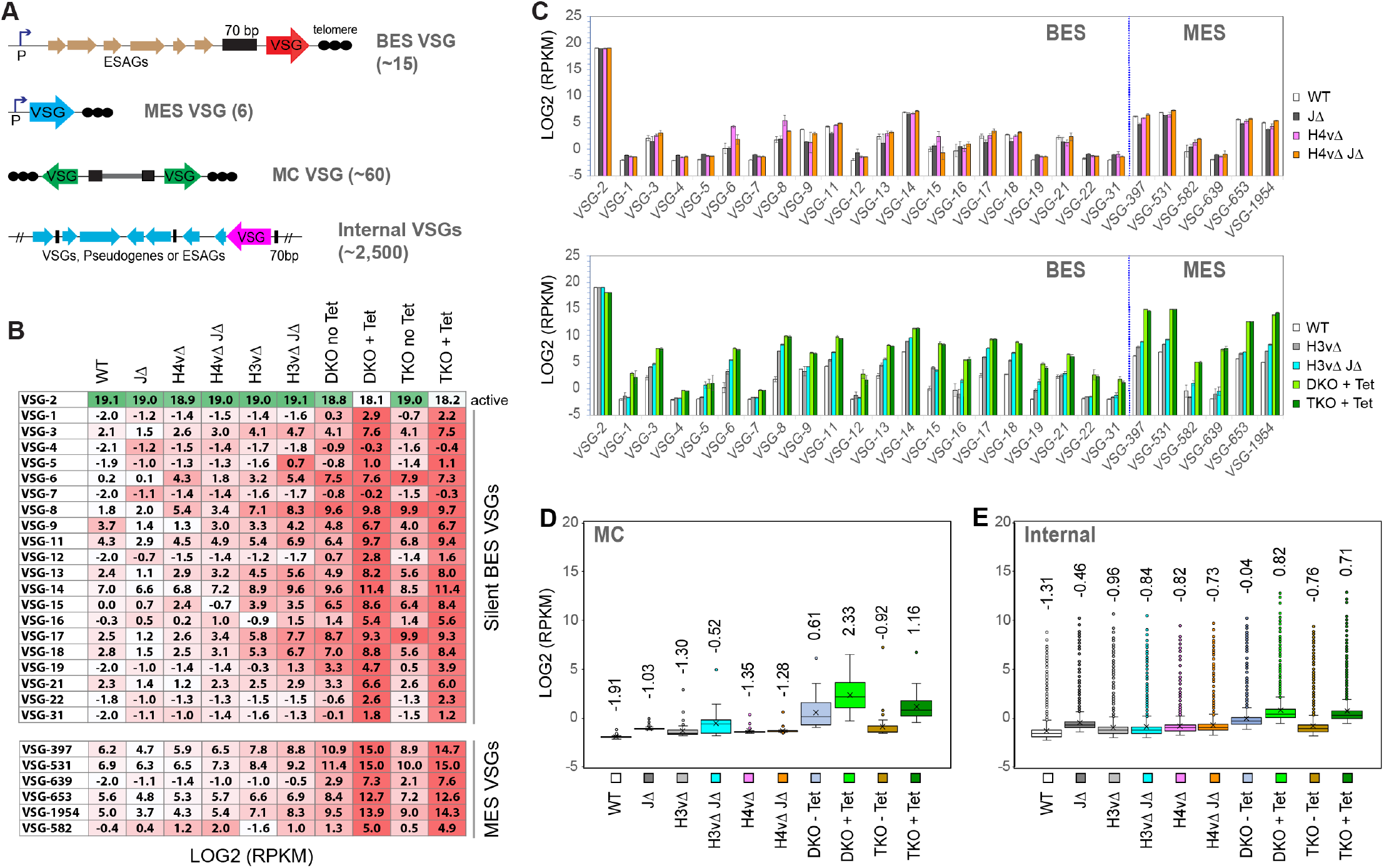
TTS-chromatin marks are requisite for VSG transcription in a synergistic manner. VSG RNA levels in WT and KO mutants were analyzed by rRNA-depleted stranded RNA-seq. RNA-seq reads were aligned to the VSGnome database^26^ and LOG2(RPKM) values were obtained for each VSG in wild type and KO mutants. **(A)** Four types of VSG loci in the genome. BES VSG (telomeric, polycistronically transcribed by RNA pol I), MES VSG (telomeric, monicistronically transcribed by RNA pol I), and MC VSG (telomeric, no promoter). Chromosome-internal VSG in arrays. **(B)** Heat map comparison between WT and KO mutants for BES and MES VSG expression levels (darker color = higher expression levels). Shown: silent VSGs (red) and active VSG2 (green). **(C)** Bar graphs comparing WT and KO mutants for BES and MES VSG expression levels. Mutants are grouped by the loss of VSG silencing phenotype: mutants that showed no change (top) and mutants that increased the expression of silent VSGs (bottom). Error bars indicate standard deviation between three replicates. **(D)** Box plot comparing WT and KO mutants for MC VSG expression levels. **(E)** Box plot comparing WT and KO mutants for chromosome internal VSG expression levels. Statistically analyses are summarized in Supplementary Table 9.

**Figure 5.**
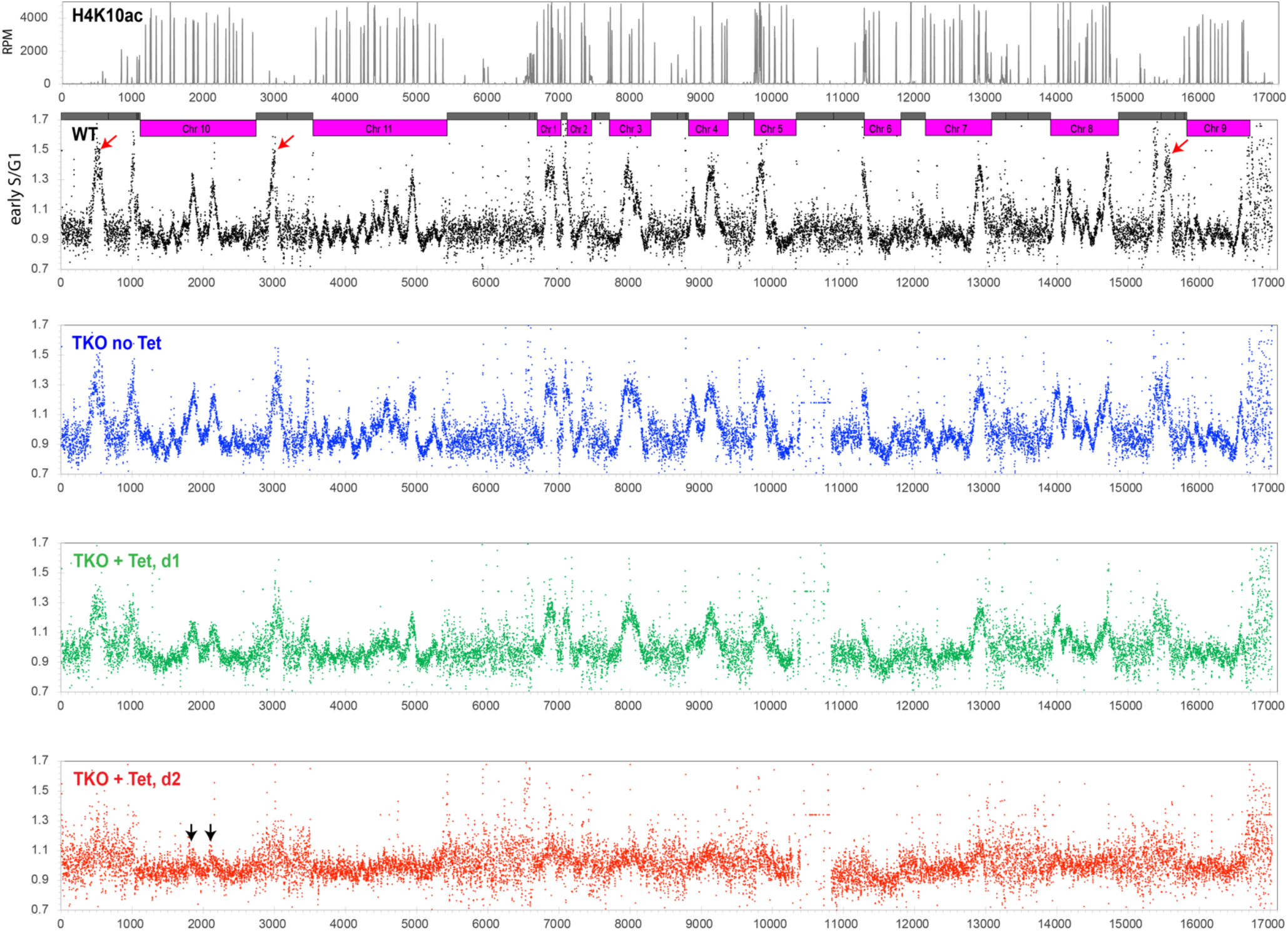
Global DNA replication impairment in TKO cells. Replication was examined by MFA-seq. TKO strain with floxed H3v-HA allele was treated with tetracycline for 0, 1 or 2 days and stained with PI. Cells in G1, early S, late S and G2 phase were FACS-sorted (Supplementary Figure 9). Genomic DNA was prepared and sequenced in the Illumina platform. Sequence reads were aligned to the Lister 427 genome and analyzed with sliding window (10kb bin, 2.5kb step). Read count ratio between early S to G1 was plotted for the whole genome, including subtelomeric regions. Shown: chromosome cores (pink) and subtelomeres (grey) in the chromosome diagram. Several of early replicating origins occur in subtelomeres of chromosome 10, 11, and 9 (red arrows). MFA-seq reads obtained from WT^32^ was re-analyzed and mapped to the Lister 427 genome for comparison. Subtelomere 6A of chromosome 6 is lost in the TKO strain.

Although MC VSGs do not have a promoter, they experience similar regulation by H3v, H4v, and J (Fig. 4D). The mean values of MC VSG transcript levels increased in mutants lacking H3v, in the order of DKO+Tet, TKO+Tet, H3vΔ JΔ and H3vΔ (high to low), though the underlying mechanism for the derepression of these promoter-less VSGs remains unclear. They may contain ‘promoter-like’ elements, which become more accessible by RNA pol I, as the heterochromatin structure is disrupted in H3vΔ H4vΔ cells. Chromosome-internal VSGs were also upregulated similarly (Fig. 4E). Statistical analyses for all VSG expression data are in Supplementary Table 9.

Müller et al observed an increased number of switched cells in H3vΔ H4vΔ cells^41^. To determine if VSG switching also increases in Tet-treated DKO and TKO cells, I examined VSG switching using flow cytometry. Because removing H3v is not reversible and Tet-treated conditional KO cells are very sick, accurately measuring the VSG switching rate posed challenges. Therefore, I looked at switching between two VSGs by staining them with antibodies conjugated with different fluorophores, the active VSG2 with Dylight 488 (green) and the silent VSG3 with Dylight 650 (red). VSG3 was chosen because VSG3 is the most frequently selected silent VSGs in switching assays in the SM strain background^19,42–44^. VSG3 expressing cells were added in to VSG2-expressing cells at 0.01%, 0.1%, 1%, and 10%, and cells were stained with anti-VSG2 and anti-VSG3 antibodies and analyzed by flow cytometry. The percent of VSG3 expressing cells mixed matched well with % VSG3 detected (Supplementary Fig. 8A, B). However, mixing of two strains expressing two different VSG coats produced a population of cells that colocalized both antibodies. This population only occurred when two strains were mixed, as they were not detected in 100% VSG2 expressing cells (VSG2 only). I speculate that there might be some cell-to-cell interaction. Using this method, silent VSG3 expression (derepression or switching) was examined in WT, DKO and TKO treated with or without Tet (Supplementary Fig. 8C). The number of double positive population increased in Tet-treated DKO and TKO cells, but I did not detect any obvious VSG3 switcher population in either sample. Because the abnormal cell population accumulated in mutants, I could not gate other VSG switchers that are not VSG3 expressors. However, at least for VSG3, it is clear that switching did not increase. I conclude that the VSG3 protein detected upon Tet induction likely arise from derepression of VSG3 locus, rather than from switching to VSG3.

### Global DNA replication impairment in TKO cells

The triple KO cells showed inefficient DNA replication and abnormal cell division, producing a large population of cells containing no or less than 2C DNA content. TSSs are also sites of *Tb*ORC1occupancy in *T. brucei*^45^, so transcription must coordinate with replication progression to prevent genome instability. To investigate global replication initiation at early replicating origins, I performed MFA-seq experiment (Marker Frequency Analysis followed by HT sequencing) in TKO cells, as previously described^32,45,46^.

PI-stained TKO cells treated with tetracycline for 0, 1, and 2 days were sorted by Fluorescence-Activated Cell Sorting (FACS) (supplementary Fig. 9). Genomic DNA was isolated from cells in G1, early S, late S, or G2 phase, and sequenced. Sequence reads were mapped to the Lister 427 reference genome and analyzed with sliding windows (10kb bin, 2.5kb step) (Fig. 5). I re-analyzed WT MFA-seq reads (which were mapped to the *Tb*927 reference genome^32^) for the Lister 427 genome. I found that subtelomeres also contained several of strong early firing origins (indicated as arrows in red in WT). Untreated TKO strain showed the same peak pattern as the WT. At day 1 post Tet induction, the peak pattern was the same but with a reduced intensity compared to day 0. Although several of small peaks were visible at some of strong origin sites at day 2 (indicated as arrows in black), no major peaks were detected, indicating a global impairment of DNA replication initiation.

Cell-cycle sorting (supplementary Fig. 9C, D) indicates that early-S phase cells should yield about 30% more DNA compared to G1 cells. While this increased DNA amount was clearly mapped to regions of early replicating origins in WT and untreated TKO cells, in TKO cells treated with Tet for 2 days, it is impossible to identify regions where this extra DNA was generated from, as no peak was detected. One possibility is that DNA synthesis may be initiated promiscuously (e.g., activating more origins including dormant ones) but replication forks progress inefficiently, producing shorter BrdU-labeled strands. Alternatively, unequal chromosome segregation in a replication-defective parent cell may produce daughter cells with one containing more DNA (early-S DNA content) and the other containing less DNA (sub-G1 DNA content).

### Emergence of surviving cell populations in cells lacking H3v and H4v

H3vΔ H4vΔ cells generated by conditional KO showed defects in growth, cell-cycle and DNA replication within two days (Fig. 1). Growth stalling occurred in some of H3vΔ H4vΔ clones obtained by transfection (Supplementary Fig. 1). These data suggest that removing H3v and H4v may cause growth arrest initially, but some cells adapt and manage to proliferate. To determine whether Tet-treated DKO cells are inviable or growth arrested, I measured the viability of DKO and TKO cells 2 days after Tet induction. The viability of Tet-treated DKO and TKO mutants was both low (Fig. 6A), indicating that H3vΔ H4vΔ is lethal primarily (94.4% and 96.9% lethal). Seven survived clones from each strain (DKO survivor, DS and TKO survivor, TS) were analyzed. They grew at different rates (supplementary Fig. 10A). DS clones showed different levels of VSG3 proteins, while TS clones showed similar levels of VSG3 protein but lower than Tet-treated TKO (Fig. 6C). Three of DS and TS clones were analyzed further in stranded RNA-seq (rRNA depleted). Three DS clones showed different patterns of BES VSG expression (Fig. 6D, left). In DS2 and DS3, silent VSG expression levels were lower than Tet-treated DKO replicates, but majority of silent VSGs in DS5 expressed the same level or more than in Tet-treated DKO cells. The level of MES VSG transcripts in DS clones was lower than Tet-treated DKO and was generally uniform. Compared to DS clones, less clonal variation occurred in TS clones in BES or MES VSG expression (Fig. 6D, right). VSG expression profiles of all replicates are summarized in box plots (Supplementary Fig. 11).

Because H4v is the main factor for transcription termination control, I did not suspect major changes in transcription profiles as H4v was already absent in DKO and TKO strains. Interestingly, transcription termination defects appear partially alleviated in DS5 (Fig. 6E, left). In addition, levels of elevated antisense transcripts in TS clones were slightly lower than Tet-treated TKO (Fig. 6E, right). Collectively, I conclude that the loss of H3v, H4v (and base J) generates toxic effects initially but cells that overcome do emerge.

**Figure 6.**
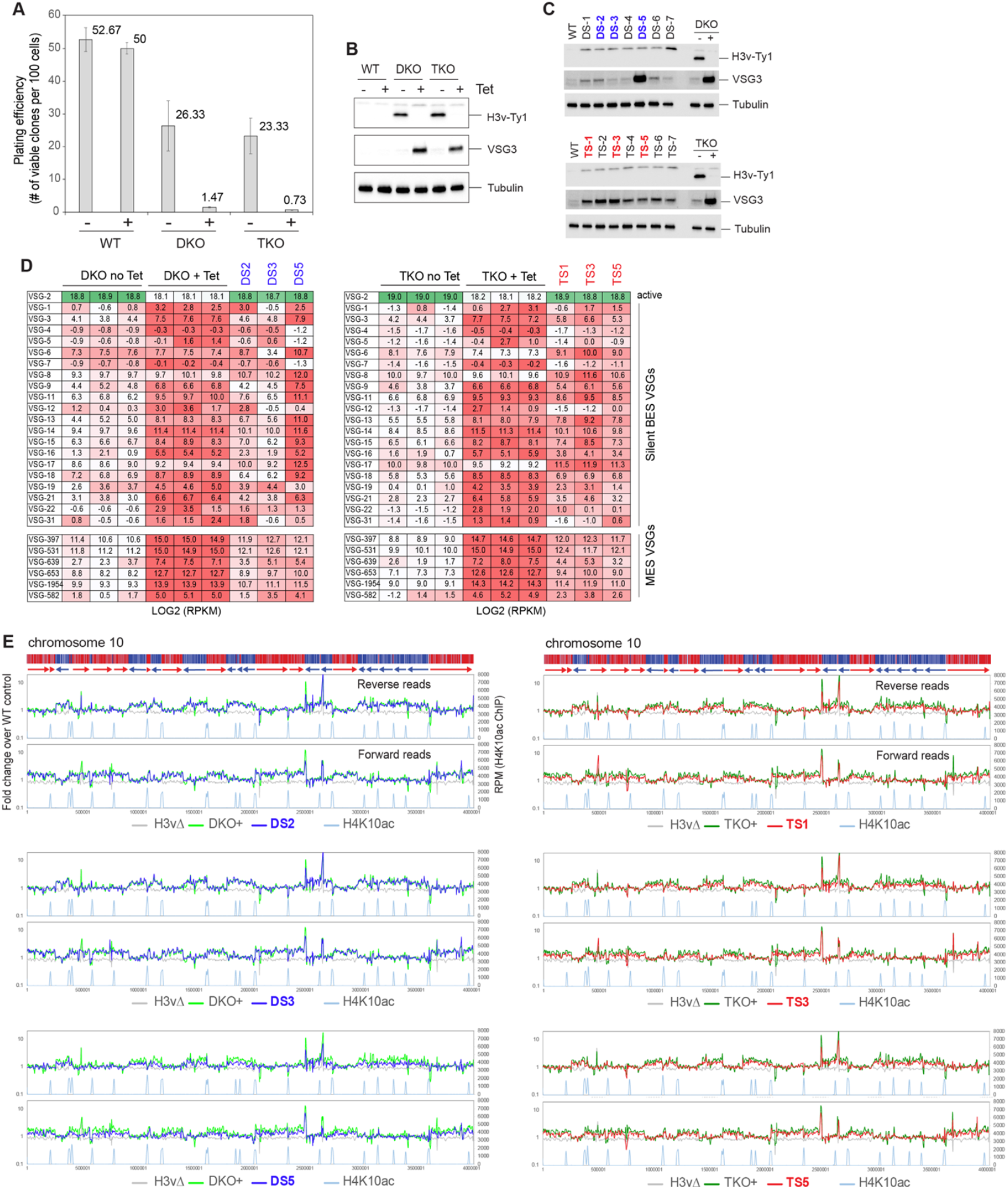
Emergence of a surviving cell population lacking H3v and H4v. **(A)** Plating efficiency after H3v-Ty1 removal in DKO and TKO strains. WT, DKO and TKO strains were treated with tetracycline for two days to remove H3v-Ty1 allele. 100 or 1000 cells were distributed in 96 well plates and the number of wells containing live cells was counted to determine the viability. Clones that are resistant to puromycin were excluded as they may grow because they still retain the H3v-Ty1-PUR allele. Error bars indicate standard deviation between three replicates. **(B)** Western blot confirming the absence of H3v-Ty1 protein in Tet-treated cells in (A). (**C)** Silent VSG3 protein expression in survivors emerged after removing H3v-Ty1. Seven of survivor clones from DKO (DS 1∼7) or TKO (TS 1∼7) were examined for silent VSG3 expression by western blot. (**D)** VSG silencing in DS and TS clones was examined by stranded RNA-seq (rRNA depletion). Heat maps compare DKO replicates (-/+ Tet) and DS clones, and TKO replicates (-/+ Tet) and TS clones for BES and MES VSG transcript levels (darker color = higher expression level). **(E)** Global transcription and antisense transcription in DS and TS clones by rRNA-depleted stranded RNA-seq (10kb bin, 2.5kb step). Fold changes compared to WT were plotted and compared with Tet-treated DKO or TKO mutant.

### H3v and H4v do not reside in the same nucleosome

To ensure that phenotypes observed from conditional KO strains were due to the H3v removal and not some artefacts, I conducted complementation experiments. I reintroduced an ‘un-floxed’ WT-H3v allele in the TKO strain (floxed H3v-HA in H4vΔ JΔ) (Fig. 7A). Cre can remove only the floxed H3v-HA allele and not the complementing WT-H3v, as shown in PCR genotyping (Fig. 7B). H3v fully complemented defects in cell growth, cell-cycle progression and VSG silencing defects (Fig. 7C-F), indicating that H3v removal caused the observed phenotypes. WT-JBP1 or JBP2 was also introduced in the TKO strain using a similar approach. Slight growth improvements were observed in JBP2 complemented Tet-treated TKO cells. JBP2 partially rescued the cell-cycle defect (Fig. 7C-E), as the number of sub-G1 cells was significantly reduced by JBP2 introduction. JBP1 also reduced the number of sub-G1 cells, but fewer than JBP2. Introduction of JBP1 or JBP2 can restore base J modification in JΔ cells at different genomic loci^7,10^. JBP1 stimulated base J modification at PTU boundaries, including TTSs and TSSs, while JBP2 did not^7^. Instead, JBP2 partially restored J modifications associated at telomere, 70bp repeats, and 177 bp repeats in minichromosomes^10^. I posit that base J (or JBP1/2 protein) is essential to coordinate DNA replication completion and cell division.

**Figure 7.**
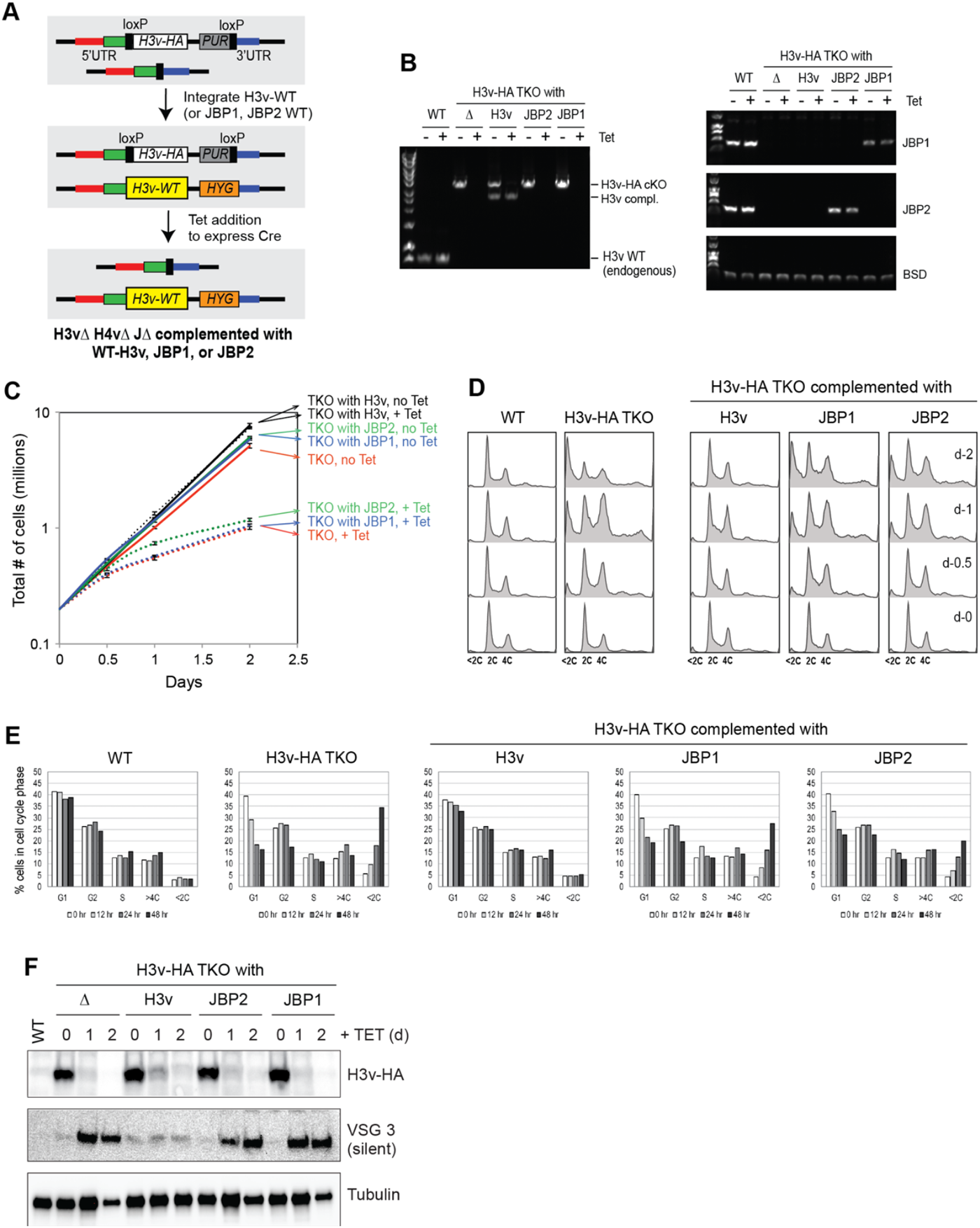
Full and partial complementation of TKO phenotypes by H3v and JBP1/2. **(A)** Strategy to generate complemented cell lines. WT-H3v, JBP1 or JBP2 gene was reintroduced to the H3v-HA TKO strain (floxed H3v-HA in H4vΔ JΔ strain) at the endogenous H3v locus where the H3v was deleted and the selection marker removed. Because the targeting vectors do not have loxP sites, Cre cannot remove the complementing gene, while it removes floxed H3v-HA allele. Resulting cells will express WT-H3v, JBP1 or JBP2 in H3vΔ H4vΔ JΔ background. **(B)** PCR genotyping confirming the removal of the floxed H3v-HA allele and presence of complemented H3v, JBP1 or JBP2 gene in Tet-untreated or Tet-treated TKO strains. **(C)** Complementation of growth defect. TKO or TKO strain integrated with WT-H3v, JBP1, or JBP2 were treated with tetracycline for 0, 0.5, 1, and 2 days as the cell count was monitored. **(D)** Complementation of cell-cycle defect. Cells treated as in (C) were fixed and stained with PI and analyzed by flow cytometry. **(E)** Graphs showing percentage of G1, G2, S, and cells with abnormal DNA content (e.g., <2C or >4C). **(F)** Complementation of silent VSG3 derepression. H3v-HA and VSG3 protein expression by western blot.

The largest subunit of Replication Protein A complex (RPA), RPA1, forms nuclear foci during replication stress and DNA damage in eukaryotes, including *T. brucei*^47–49^. Tet-treated TKO cells displayed nuclear *Tb*RPA1 foci (Supplementary Fig. 12). Collectively, the data suggest that the loss of H3v and H4v induces replication stress and activates the DNA damage response (DDR) including *Tb*RPA1 recruitment to the lesions, and cell-cycle arrest in a J dependent manner.

The genetic interaction thus far strongly suggests that H3v and H4v do not reside in the same nucleosome and this was confirmed by co-immunoprecipitation (co-IP) assays using H3v-HA and/or Flag-H4v expressing strains (Supplementary Fig. 13).

## Discussion

The data here indicate that genome-wide transcription termination readthrough neither interferes with DNA replication nor with cell growth in *T. brucei*. H3v and H4v deletion led to problems in DNA synthesis and growth in DKO and TKO mutant strains. However, Tet-treated DKO and TKO cells produced about 5.6% and 3.1% viable clones. Since the high rate suggested another mechanism than genetic suppression, I hypothesized that epigenetic adaptation may underlie this effect. Initial growth defects in Tet-treated DKO and TKO cells could be caused due to unprotection of TTS DNA. In mice, cells with less nucleosomes accumulated more DNA damage and nucleosome-occupied regions had more protection from irradiation induced DNA damage^50^. Therefore, nucleosome-free TTS DNA may become fragile, accumulate DNA lesions, and activate DNA damage response in Tet-treated DKO and TKO cells. To protect TTS DNA, cells may attempt to randomly deposit core H3-H4, perhaps with specific PTMs at TTSs, most of which are likely unsuccessful. If a cell deposited a functional pair of H3-H4 PTMs capable of stable nucleosome formation at TTSs, these cells could emerge and survive (Fig. 8A). Several of H3v and H4v PTM are now identified^8^, although their functions are not yet known, as well as chromatin proteins associated at TTSs, which include readers and writers of histone PTMs (BDF7, PHD2, PHD4, DOT1A)^51^. H4 (or H3) with a specific PTM may associate with H3v (or H4v) for depositing to TTSs. I conclude that PTMs of histones and histone variants marking TTSs are critical for regulating transcription termination.

**Figure 8.**
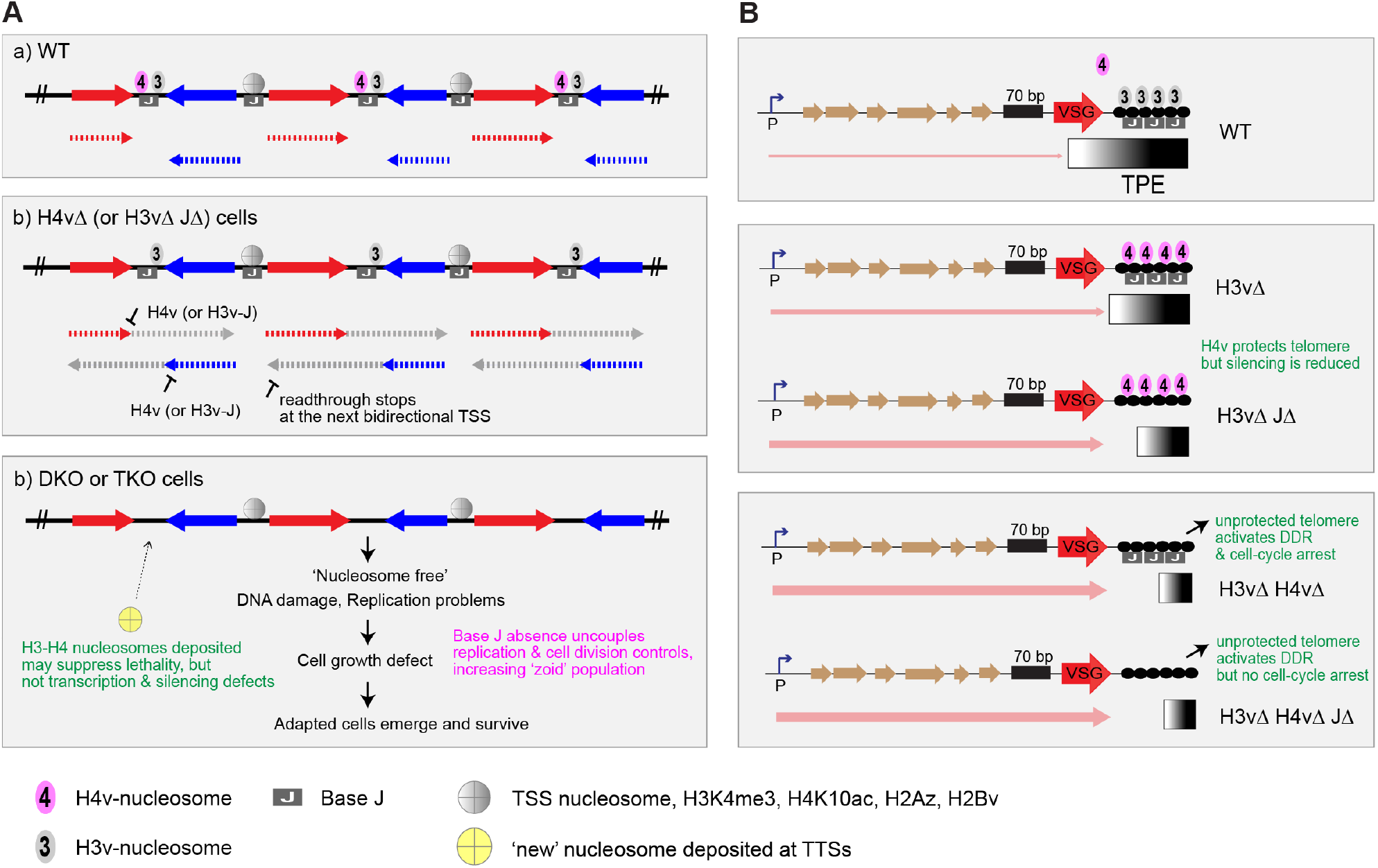
Roles of H3v, H4v and base J in transcription and genome maintenance. **(A)** At TTS, J-modified DNA wraps around nucleosomes containing H3v or H4v. H4v nucleosomes signal the RNA pol II machinery to stop at TTSs. Pol II that missed the stop sign continues to progress until it reaches the next divergent TSS. Simultaneous absence of H3v and J affects H4v nucleosome function by changing the TTS chromatin structure. Upon removal of H3v in DKO or TKO cells, nucleosome-free TTS DNA becomes fragile, accumulates replication stress and DNA lesions, and activates DDR. When J is present, nucleosome-free TTSs may signal cells to slow down replication and cell-cycle progression to repair the damage. However, TKO cells without J go through mitosis and cytokinesis with incomplete replication and unrepaired DNA, producing daughter cells lacking DNA. A cell may deposit a functional pair of H3-H4 PTMs. If a survivor clone with new epigenetic marks can emerge, it could form stable nucleosomes and TTS chromatin structure and inherit. **(B)** The synergistic effect caused by H3v and H4v double KO in VSG silencing suggests that H4v nucleosome may bind telomere repeats in the absence of H3v. The loss of both H3v and H4v nucleosomes at telomeric regions could leave telomeres unprotected, leading to chromosome abnormalities, disruption of heterochromatin structure and loss of VSG silencing. Differing degrees of VSG derepression level between BES VSGs in a DS clone and also between DS clones suggests that individual telomere in survivors may not have the same epigenetic code. Partial complementation of cell-cycle defect by JBP1 or 2 suggests that JBP1/2 or base J at TTS and/or telomere may coordinate the completion of replication with cell-cycle progression.

Sub-G1 population increase was more pronounced in Tet-treated TKO, than DKO, indicating that J has a special function in cell-cycle control. In other eukaryotes, activation of cell-cycle checkpoint upon DNA damage or replication stress leads to cell-cycle arrest at G1 or G2/M, or delays in S-phase progression^52^. Thus, accumulation of sub-G1 population in *T. brucei* is an unusual phenotype for a checkpoint defective mutant. Sub-G1 populations can arise through an unequal segregation of chromosome and/or uncoupling of mitosis/cytokinesis and replication completion. In the presence of base J, nucleosome-free TTSs may signal cells to slow down replication and cell-cycle progression to repair the damage. When base J is absent, H3vΔ H4vΔ cells may fail to respond to this signal and proceed with mitosis and cytokinesis with incomplete replication and unrepaired DNA, produce daughter cells lacking DNA and induce cell death.

*T. brucei* JBP proteins prompted the discovery of mammalian TET enzymes (Ten Eleven Translocation) that convert 5-methylcytosine (5mC) to hydroxy-5mC (5hmC), similar to JBP proteins converting methyl-U (T) to hmU. TET enzymes are important factors for reprogramming during development and differentiation^53,54^. TET1/2 depleted trophoblast cells showed increased polyploidy due to endoreduplication, defective G2-M transition and centriole separation^55^. Regions adjacent to DNA damage accumulate 5hmC modifications in a TET2 dependent manner in human cells. Mouse ESC lacking TET1-3 enzymes exhibited abnormal chromosome segregation that was further aggravated by aphidicolin induced replication stress^56^. Here, data suggest that base J (JBP1/2) may bridge between chromosome abnormalities and cell-cycle control, which was unnoticeable under normal conditions. Although not identical, the experiments described here demonstrate shared roles between epigenetic modifiers from evolutionarily distant organisms.

H3v, but not H4v, bind telomere repeats^5^. Consistent with this, deletion of H3v, but not H4v, disrupts VSG silencing. If canonical H3-H4 nucleosomes occupy telomeres in H3vΔ mutant, the same VSG silencing phenotype should occur in the H3vΔ and H4vΔ H3vΔ mutants. However, the synergistic VSG silencing defect observed in Tet-treated DKO indicates that H3vΔ and Tet-treated DKO cells may have different telomeric chromatin structure. One possibility is that H4v nucleosomes occupy telomeres in the H3vΔ mutant. In Tet-treated DKO cells, telomeres may be nucleosome free or occupied by H3-H4 nucleosomes that cannot maintain telomeric heterochromatin. Telomere unprotection and heterochromatin structure disturbance could lead to the loss of telomere silencing and activation of DDR (Fig. 8B). JBP2 expression in JΔ cells partially restored base J levels at telomere repeats^7,10^, and I found that JBP2 can partially rescue cell-cycle defects in the Tet-treated TKO cells. JBP2 or telomeric J may play critical roles in replication stress and DNA damage induced cell-cycle checkpoint control in *T. brucei*.

Upregulation of antisense transcription also occurs in H4vΔ and Tet-treated TKO mutants, but the increase in polyadenylated antisense RNA only occurred in Tet-treated TKO cells. Polyadenylation machinery may interact with TTS chromatin to distinguish coding region from non-coding within precursor RNAs, preventing polyadenylation of antisense transcripts. TFIIS2-2 (transcription elongation factor) is enriched at TTSs and interacts with PAF1 complex in *T. brucei*^51^. PAF1 contributes to chromatin remodeling, transcription elongation and polyadenylation in yeast and mouse cells^57,58^. Interestingly, PAF1 complex interacts with JBP3 in *Leishmania tarentolae*^59^. JBP3 is a J-binding protein 3 recently identified in *T. brucei* and *L. tarentolae*. JBP3 depletion also led to transcription readthrough at TTSs in *T. brucei*^59,60^. In plant, polyadenylation of antisense transcripts affects sense transcript level by interacting with RNA-binding protein, RNA 3’ processing factor, histone H3 methyltransferase and demethylase^61^. Future studies will elucidate whether polyadenylated antisense transcripts in the triple KO modulate cell viability and how TTS chromatin proteins contribute to polyadenylation regulation.

## Materials and Methods

### *Trypanosoma brucei* strains and plasmids

Bloodstream form *Trypanosoma brucei* (the Lister 427 antigenic type MITat1.2 clone 221a) was cultured in HMI-9 at 37 °C^62^. All cell lines were constructed in a ‘Single Marker (SM)’ background, which has a tetracycline-inducible system constitutively expressing T7 RNA polymerase and Tet repressor^63^. CRE gene under Tet Operator control is stably integrated at an rDNA spacer locus and its expression can be induced by tetracycline addition.

Genes encoding JBP1, JBP2, H3v, and H4v were deleted using knockout (KO) vectors. These vectors have gene-specific homology sequence arms that are homologous to upstream or downstream sequences of a target gene. Between upstream and downstream homology arms, a selection marker flanked by loxP sites is located. A positive selection marker (hygromycin-resistance gene, HYG, or puromycin-resistance gene, PUR) conjugated with *Herpes Simplex* Virus Thymidine Kinase gene (HSVTK or TK) was used. After deleting both alleles of a gene, tetracycline (Sigma-Aldrich) can be added to express Cre to remove the floxed HYG-TK and PUR-TK markers, so these selection markers can be reused to delete the next gene. To generate a conditional null mutant, a ‘conditional KO (cKO or floxed)’ allele was introduced using a cKO vector, which is the same as a KO vector except that H3v-HA, H3v-Ty1, or Flag-H4v gene is inserted in between of the 5’ loxP and a selection marker gene. Cre can remove a region between two loxP sites (including the H3v-HA or Flag-H4v allele), thereby generating cells that no longer express H3v or H4v gene upon tetracycline addition. WT (HSTB-904), JΔ (HSTB-778), H3vΔ (HSTB-881), and H3vΔ JΔ (HSTB-868) were generated previously^19^. In this study, following strains were generated.

H4vΔ strain: One H4v allele in the wild type strain (HSTB-904) was deleted with pKP26 vector containing a H4v-KO-HYG-TK cassette (‘H4v-KO’ indicates that the cassette contains homology arms for H4v deletion), creating a H4vΔ/+ heterozygote (HSTB-1039). The second H4v allele in HSTB-1039 was deleted with pKP27 containing a H4v-KO-PUR-TK, creating a H4vΔ/Δ homozygote (HSTB-1051). Selection markers in HSTB-1051 were removed by Cre expression (HSTB-1067).

H4vΔ JΔ strain: One H4v allele in the JΔ strain (HSTB-778) was deleted with pKP26 (H4v-KO-HYG-TK), creating a H4vΔ/+ JΔ (HSTB-1041). The second H4v allele in HSTB-1041 was deleted with pKP27 (H4v-KO-PUR-TK), creating a H4vΔ/Δ JΔ (HSTB-1055). Selection markers in HSTB-1055 were removed by Cre expression (HSTB-1069).

H4vΔ H3vΔ strain: One H4v allele in the H3vΔ strain (HSTB-881) was deleted with pKP26 (H4v-KO-HYG-TK), creating a H4vΔ/+ H3vΔ (HSTB-1043). The second H4v allele in HSTB-1043 was deleted with pKP27 (H4v-KO-PUR-TK), creating H4vΔ/Δ H3vΔ strains (HSTB-1260, 1261, 1262, 1263, 1264, and 1265).

An attempt to generate H4vΔ H3vΔ JΔ strain: One H4v allele in the H3vΔ JΔ strain (HSTB-868) was deleted with pKP26 (H4v-KO-HYG-TK), creating H4vΔ/+ H3vΔ JΔ (HSTB-1045). pKP27 (H4v-KO-PUR-TK) was transfected in HSTB-1045. Clones were obtained initially but did not expand further.

Floxed H3v-Ty1 in H4vΔ strain (DKO with floxed H3v-Ty1): One allele of H3vΔ was deleted in the H4vΔ strain (HSTB-1067) with pDS88 (H3v-KO-HYG-TK), creating a H3vΔ/+ H4vΔ strain (HSTB-1074). Cre was induced to remove a HYG-TK selection marker (HSTB-1078). The remaining H3v allele in the H3vΔ/+ H4vΔ strain (HSTB-1078) was replaced with the floxed H3v-Ty1-PUR (pKP33 vector), creating a floxed H3v-Ty1/Δ H4vΔ strain (HSTB-1082, DKO). H3v-Ty1 allele can be removed by tetracycline addition, generating H3vΔ H4vΔ double KO cells (Tet-treated DKO).

Floxed H3v-Ty1 in H4vΔ JΔ strain (TKO with floxed H3v-Ty1): One allele of H3vΔ was deleted in the H4vΔ JΔ strain (HSTB-1069) with pDS88 (H3v-KO-HYG-TK), creating a H3vΔ/+ H4vΔ JΔ strain (HSTB-1076). Cre was induced to remove a HYG-TK selection marker (HSTB-1080). The remaining H3v allele in the H3vΔ/+ H4vΔ JΔ strain (HSTB-1080) was replaced with the floxed H3v-Ty1-PUR (pKP33 vector), creating a floxed H3v-Ty1/Δ H4vΔ JΔ strain (HSTB-1088, TKO). H3v-Ty1 allele can be removed by tetracycline addition, generating H3vΔ H4vΔ JΔ triple KO cells (Tet-treated TKO).

Floxed H3v-HA in H4vΔ JΔ strain (TKO with floxed H3v-HA): The remaining H3v allele in the H3vΔ/+ H4vΔ JΔ strain (HSTB-1080) was replaced with floxed H3v-HA-PUR-TK (pDS84 vector), creating a floxed H3v-HA/Δ H4vΔ JΔ strain (HSTB-1142). H3v-HA allele can be removed by tetracycline addition, generating H3vΔ H4vΔ JΔ triple KO cells.

Floxed Flag-H4v in H3vΔ JΔ strain (TKO with floxed Flag-H4v): The remaining wild type H4v allele in the H4vΔ/+ H3vΔ JΔ strain (HSTB-1045) was replaced with a floxed Flag-H4v-PUR-TK (pKP49), creating a floxed Flag-H4v/Δ H3vΔ JΔ strain (HSTB-1098). Flag-H4v allele can be removed by tetracycline addition, generating H4vΔ H3vΔ JΔ triple KO cells.

Floxed H3v-HA and floxed Flag-H4v in JΔ strain (TKO with floxed H3v-HA and floxed Flag-H4v): Floxed Flag-H4v construct was transfected in floxed H3v-HA/Δ H4vΔ/Δ and JΔ strain (HSTB-1142), creating a strain with floxed H3v-HA/Δ floxed Flag-H4v/Δ JΔ (HSTB-1246).

TKO strain expressing HA-tagged TbRPA1: One allele of RPA1 was epitope tagged with 3xHA by one-step PCR integration using a pMOTag-3H vector^64^, in the floxed H3v-Ty1 H4vΔ JΔ (HSTB-1088), generating HSTB-1093.

Bloodstream form *T. brucei* strains were maintained in HMI-9 media containing necessary antibiotics at the following concentrations: 2.5 µg/ml of G418 (Sigma-Aldrich); 5 µg/ml blasticidin (InvivoGen); 1 µg/ml phleomycin (InvivoGen); 5 µg/ml hygromycin (InvivoGen); 0.1 µg/ml puromycin (InvivoGen); 1 µg/ml tetracycline (Sigma-Aldrich); 35µg/ml ganciclorvir (Sigma-Aldrich). *T. brucei* strains, plasmids, and oligonucleotides used in this study are listed in Supplementary Table 1.

### Western blot

5 to 10 million cells were collected and suspended in the Laemmli buffer and separated on an SDS-PAGE gel (Bio-RAD). Following transfer to nitrocellulose membrane (GE Healthcare), the proteins were analyzed using mouse anti-Ty1, mouse anti-HA (Sigma-Aldrich), mouse-anti-Flag (Sigma-Aldrich), mouse anti-VSG3 (Antibody & Bioresource Core Facility, MSKCC), rabbit anti-VSG3, rabbit H3 (Abcam), or mouse anti-tubulin^65^ antibodies.

### Flow cytometry analysis for cell-cycle, BrdU pulse, and VSG switching experiments

About 10 million cells of WT, DKO and TKO strain treated or untreated with tetracycline were collected and fixed with ice-cold 70% ethanol. To stain DNA, fixed cells were incubated with 50 µg/ml Propidium Iodide (PI, Sigma-Aldrich) and 200 µg/ml RNase A (Sigma-Aldrich) in PBS (Corning) at 37 °C for 30 min. Cell-cycle progression was analyzed using the LSR II or Via (BD Biosciences), FACSdiva software (BD Biosciences), and FlowJo software package (FlowJo).

BrdU pulse experiments were performed as described previously^66^, with some modifications^32^. BrdU (Sigma-Aldrich) was added to a final concentration of 500 µM to cell cultures and incubated for 40 min. Cells were then fixed with ice-cold 70% ethanol, incubated in 0.1N HCl (Fisher Scientific) with 0.5% Triton X-100 (Fisher Scientific) and washed with PBS containing 0.5% Tween 20 and 1% BSA (Jackson Immunoresearch). To detect BrdU incorporation, fixed cells were incubated with 1 µg/ml mouse anti-BrdU antibody (Fisher Scientific; BD Pharmingen) at room temperature for 2 hours and incubated with donkey anti-mouse-Alexa 488 (Invitrogen Molecular Probes) at 4 µg/ml for 45 min at room temperature in the dark. Cells were incubated in PBS containing 5 µg/ml PI and 250 µg/ml RNase A for 30 min at room temperature in the dark, and then analyzed by flow cytometry using LSR II or Via (BD Biosciences), FACSdiva software (BD Biosciences), and FlowJo software package (FlowJo).

VSG switching from VSG2 to VSG3 was examined by staining cells with antibodies conjugated with different fluorophores. VSG2 expressing cells containing 0%, 0.01%, 0.1%, 1%, or 10% of VSG3 expressing cells, and Tet-untreated or treated WT, DKO and TKO cells were prepared. 6 million live cells were stained with mouse anti-VSG2 antibody (Antibody & Bioresource Core Facility, MSKCC) conjugated with Dylight 488 (Abcam) and mouse anti-VSG3 antibody (Antibody & Bioresource Core Facility, MSKCC) conjugated with Dylight 650 (Abcam). One million cells were analyzed with Via (BD Biosciences).

### Stranded RNA-seq with rRNA depletion

WT, KO mutants, Tet-untreated DKO and TKO, and Tet-treated (for 2 day) DKO and TKO cells in triplicate, three DS clones and three TS clones were prepared. About 50 million cells were collected and total RNA was extracted using the RNA Stat-60 (Tel-Test) according to the manufacturer’s protocol and quantified on a NanoDrop2000c and further cleaned using RNeasy kit (Qiagen). rRNA was removed using the Ribo-Zero kit (Illumina) and stranded RNA-seq libraries were prepared using random hexamer and NEB directional RNA library prep kit (NEB) and sequenced on the NovaSeq 6000 PE150.

Read quality was analyzed using the FastQC program and reads were trimmed using the Trim Galore program from Galaxy (https://usegalaxy.org) (parameters: phred 20, stringency 1bp, error 0.1, length 20) and aligned with Bowtie 2^33^ to the Lister 427^41^, *Tb*927v5 (www.tritrypdb.org), or VSGnome^26^, and analyzed using SeqMonk algorithm (http://www.bioinformatics.babraham.ac.uk/projects/seqmonk/) from Babraham Bioinformatics.

To examine transcription profiles at chromosome level, each chromosome was binned at 5 kb resolution with 1 kb step (or 10kb bin, 2.5kb step for DS and TS clone analyses) as described previously^32^. Read Per Million mapped reads (RPM) values from forward or reverse reads only were generated (Supplementary Table 5, Table 11) and fold changes between WT and mutant were analyzed. To compare gene expression between WT and mutants, reads mapping to about 8,428 CDSs were analyzed. Read Per Kilobase per Million mapped reads (RPKM) values were generated from all reads, reads mapping to opposite direction of genes (sense transcription) or reads mapping the same direction as genes (antisense transcription) (Supplementary Table 6). Box plots were generated. To examine VSG expression, trimmed reads were aligned to the VSGnome (retrieved from http://tryps.rockefeller.edu)^26^. RPKM values (Supplementary Table 9) were displayed in box plots. Alignment report, correlation between replicates, and PCA analysis are summarized in Supplementary Table 2-4. Statistical analyses for box plots are summarized in Supplementary Table 7 and 9.

### Stranded RNA-seq with poly A selection

WT, JΔ, H4vΔ, H4vΔ JΔ, and Tet-treated or untreated TKO strains were grown in triplicate. About 50 million cells were collected and total RNA was extracted using the RNA Stat-60 (Tel-Test) according to the manufacturer’s protocol and quantified on a NanoDrop2000c and further cleaned using RNeasy kit (Qiagen). RNA-seq libraries were prepared from 500ng of RNA samples using an oligo dT based method with 15 cycles of PCR amplification with the Illumina TruSeq mRNA stranded kit (Illumina) and sequenced on the Illumina HiSeq 2000 v4 (50-bp single end read). Reads were analyzed for transcription profile (Supplementary Table 8) and gene expression as above (Supplementary Table 7). Alignment report, correlation between replicates, and PCA analysis are summarized in Supplementary Table 2-4. Statistical analyses for box plots are summarized in Supplementary Table 7.

### MFA-seq

MFA-seq was performed as reported previously^45^, with some modifications^32^. Floxed H3v-HA TKO cells (HSTB-1142) were treated with tetracycline for 0, 1, and 2 days. About 100 million cells at ∼1×10^6^/ml density were fixed with ice-cold 70% ethanol and DNA stained with 50 µg/ml PI in PBS. Cells in G1, early S, late S and G2 phases were FACS-sorted (BD Aria3: BD Biosciences) and genomic DNA was isolated using the QIAamp DNA Blood mini kit (Qiagen). About 10 ng of genomic DNA was fragmented using the NEBNext dsDNA fragmentase (NEB) and sequencing libraries were generated using the NEBNext Ultra DNA Library prep kit (NEB) and NEB multiplex oligos (NEB) for Illumina according to the manufacturer’s protocol. Sequencing was performed on an Illumina HiSeq 2000 sequencer (50-bp single end read). Read quality was analyzed as above; quality checked using FastQC, trimmed using Trim galore, aligned with Bowtie 2 to the Lister 427 genome. About 3 ∼ 15 million reads were analyzed using the SeqMonk algorithm. To examine replication initiation profiles, chromosomes were binned at 10 kb resolution with 2.5 kb step and RPM values were generated with SeqMonk (Supplementary Table 10). RPM values from the S phase samples were normalized to values from G1 phase and plotted.

### RPA1 immunoflueorescence

Floxed H3v-Ty1 TKO strain expressing *Tb*RPA1 tagged with HA (HSTB-1093) was treated with tetracycline for 0, 1, and 2 days. Cells were fixed with 0.5% paraformaldehyde (Thermo Fisher Scientific, Pierce) for 10 minutes, permeabilized with 0.2% NP-40 (Sigma-Aldrich) in PBS, incubated with mouse anti-HA antibody and then with the secondary antibody conjugated with Donkey-anti-mouse Alexa 567 (Invitrogen Molecular Probes). DNA was stained with 0.5 mg/ml DAPI (Sigma-Aldrich). Images were captured using a Zeiss Axioplan 2 fluorescence microscope and edited with Adobe Photoshop.

### Co-immunoprecipitation and western blot

Nuclear extract was prepared from about 10^8^ cells using Chromatin Extraction kit (Abcam). Extracts were immunoprecipitated with rabbit anti-HA (Sigma) or rabbit anti-Flag (Sigma) antibodies and precipitated proteins were analyzed by western blot using mouse anti-HA (Sigma) or mouse anti-Flag (Sigma), rabbit anti-H3 antibodies (Abcam).

## Acknowledgements

This work was supported by the National Institute of Allergy and Infectious Diseases of the National Institutes of Health (grant number R01AI127652 to H.K.). I would like to thank Svetlana Mazel and Songyan Han (Rockefeller University, Flow Cytometry Core) for help with cell sorting; Chingwen Yang (Rockefeller University, Genomics Core) for help with MFA-seq; Dewi Harjanto (Rockefeller University and Neon Labs) for discussions with regard to bioinformatics analysis tools; Nina Papavasiliou & Esteban Erben (DKFZ, Germany), and Genomics & Proteomics Core Facility at DKFZ for help with stranded RNA-seq (poly A); Novogene for NGS library prep and rRNA depleted stranded RNA-seq (Beijing, China): and Brandi Mattson at Life Science Editors for help with manuscript editing.

**Supplementary Information** is available online.

## Data availability

MFA-seq and stranded RNA-seq raw sequence files have been deposited to the Sequence Read Archive (accession number PRJNA727846).

