## supplementary figures for "Site-specific epigenetic marks in *Trypanosoma brucei* transcription termination, antigenic variation, and proliferation"

### Supplementary Information

#### Supplementary Figure 1

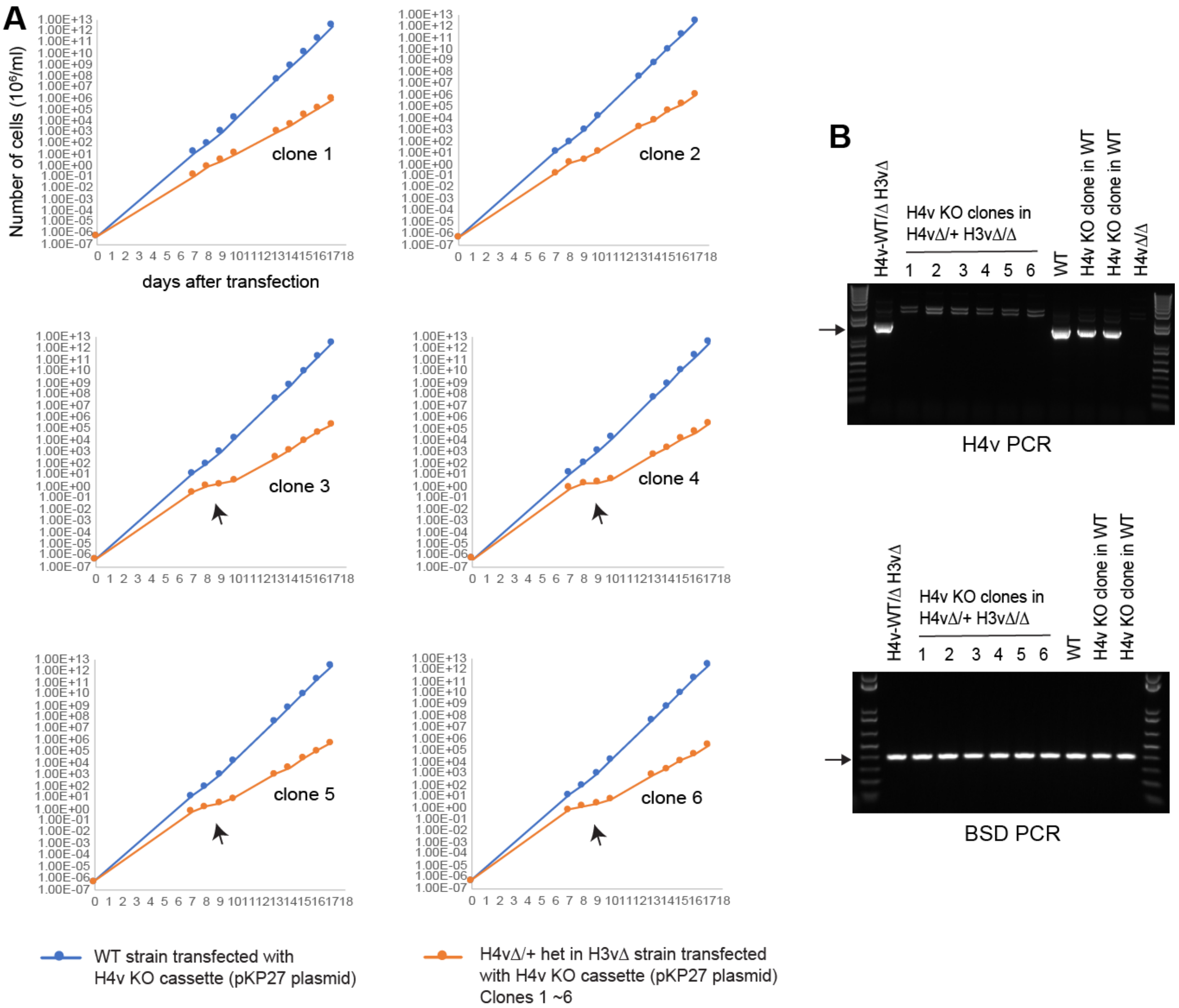


**Supplementary Figure 1. Growth of H3vΔ H4vΔ cell lines generated by stable transfection of a H4v KO cassette (linearized pKP27 plasmid) in H4vΔ/+ heterozygote H3vΔ strain. (A)** Cell growth: WT and a H4vΔ strain with one H3v allele deleted (H3vΔ/+ het in H4vΔ) were transfected with linearized pKP27 (H4v KO vector with PUR-TK marker). Clones were monitored for cell growth for 17 days after the day of transfection. **(B)** PCR genotyping confirming the absence of H4v. BSD PCR is shown as a control.

#### Supplementary Figure 2

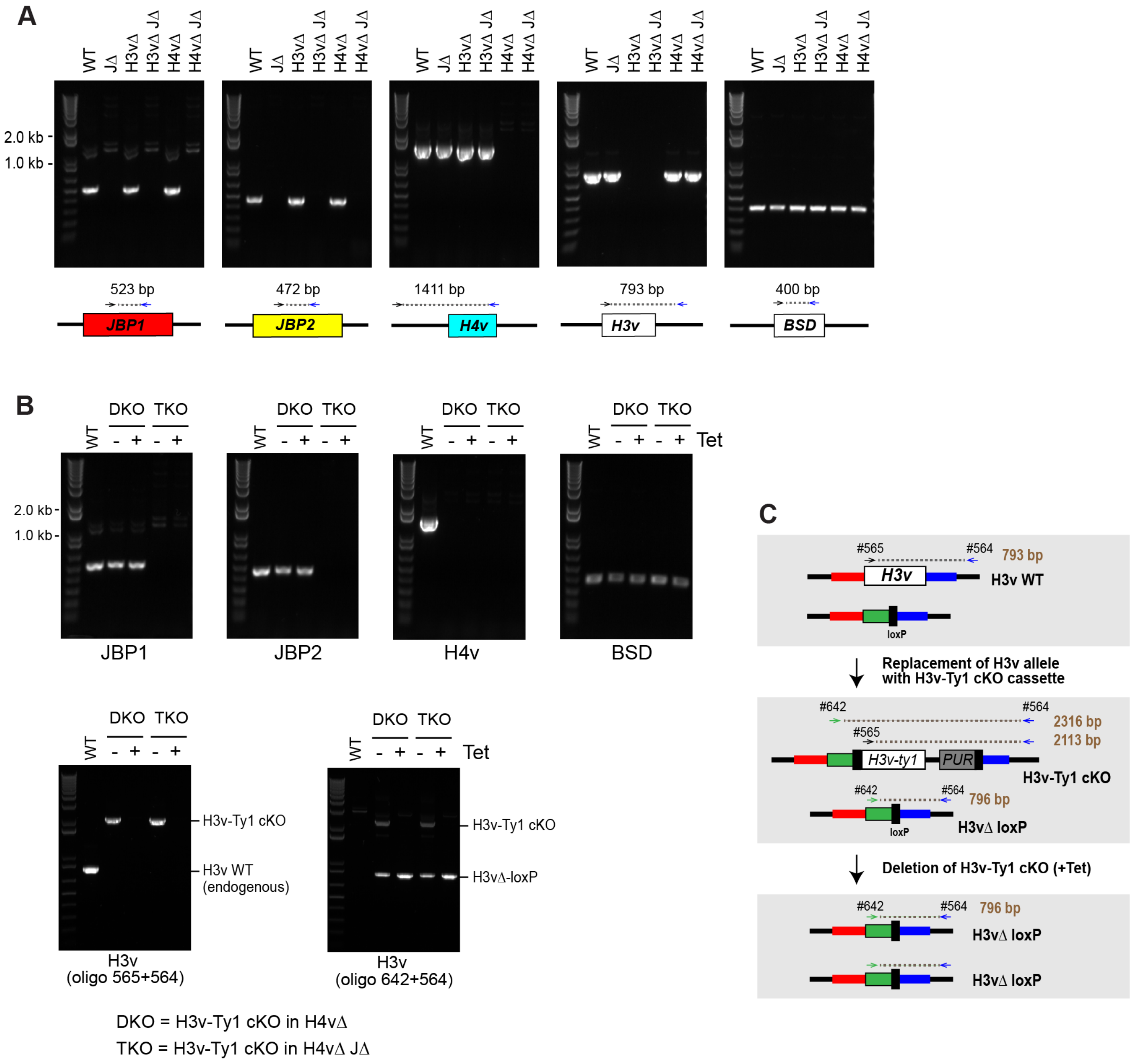


**Supplementary Figure 2. PCR genotyping of KO mutants and conditional KO mutants (DKO and TKO). (A)** PCR genotyping confirms presence or absence of JBP1, JBP2, H3v, and/or H4v in KO mutants **(B)** PCR genotyping confirms the absence of the floxed H3v-Ty1 allele in tetracycline treated DKO and TKO strains. H4v, JBP1 and JBP2 were also PCR genotyped in these strains. **(C)** Diagram showing PCR genotyping strategy for conditional H3v-Ty1 KO.

#### Supplementary Figure 3

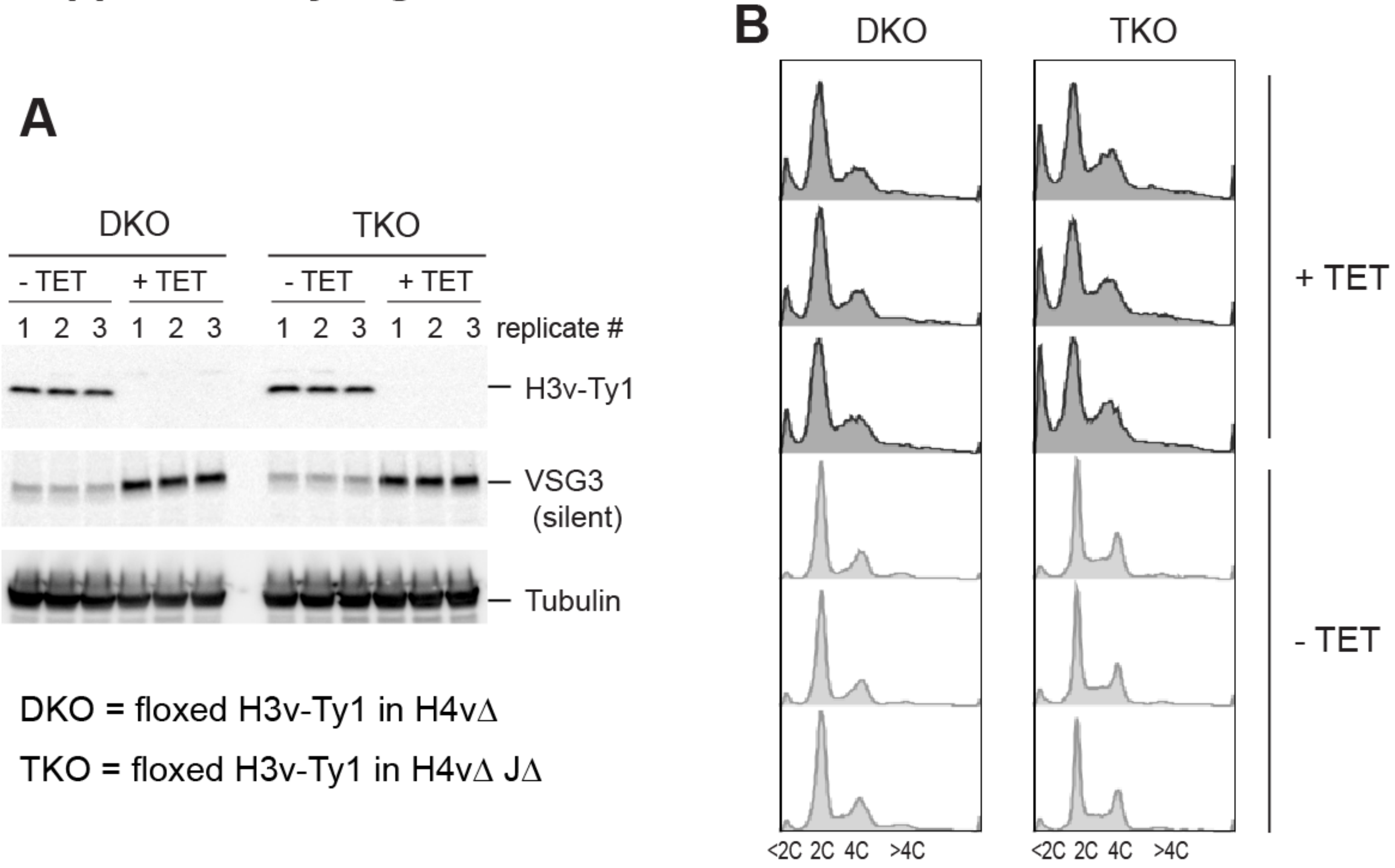


**Supplementary Figure 3. Confirmation of H3v-Ty1 removal in DKO and TKO strains with or without tetracycline in triplicate (samples used in rRNA-depleted stranded RNA-seq). (A)** Removal of H3v-Ty1 protein in Tet-treated DKO and TKO cells by western blot. **(B)** Cell-cycle profile by flow cytometry

#### Supplementary Figure 4

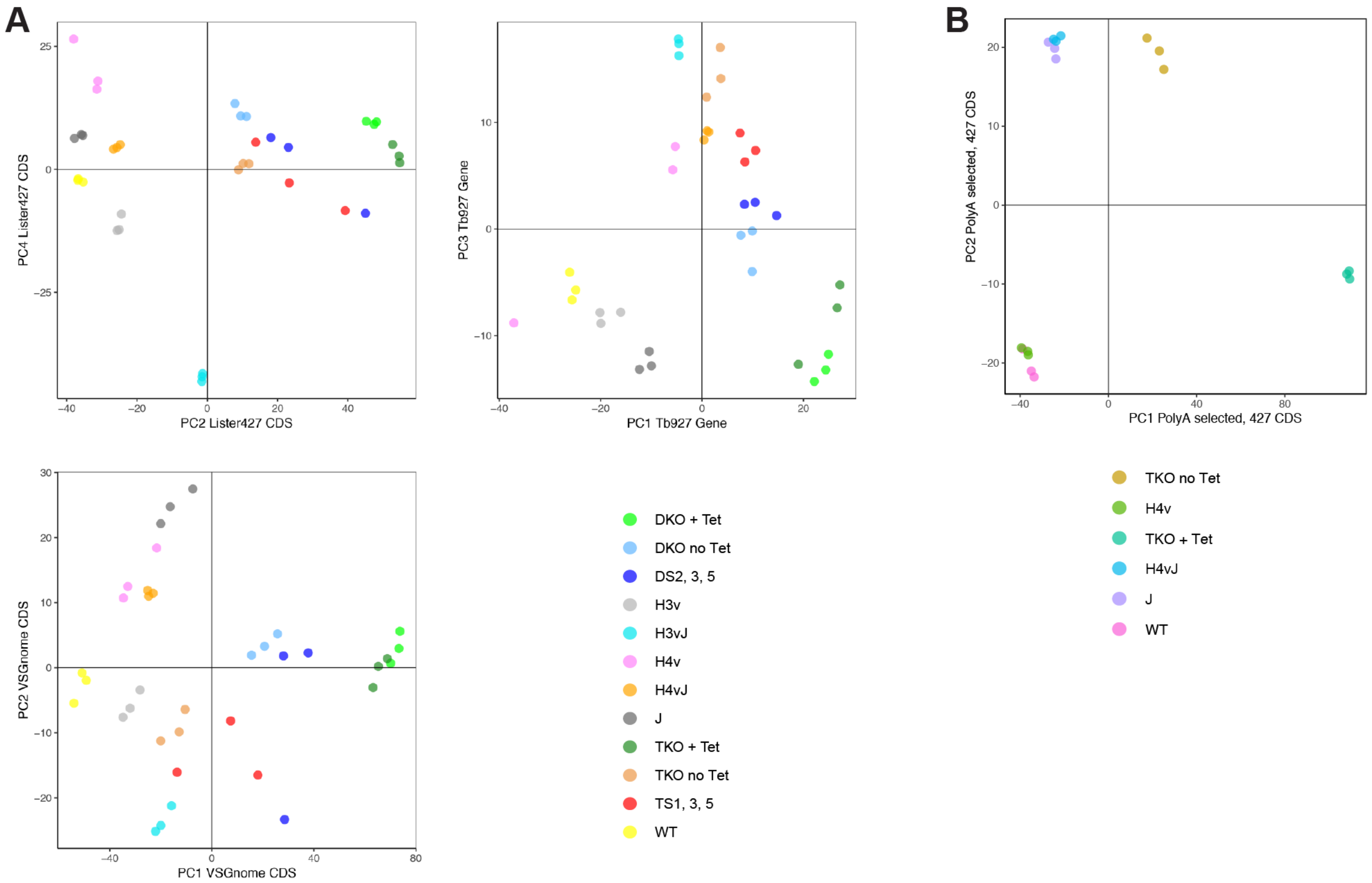


**Supplementary Figure 4. (A)** PCA plots obtained from triplicated samples used for rRNA-depleted stranded RNA-seq. Sequence reads were mapped to CDSs in the Lister 427 genome, genes in the *Tb*927 v5 genome, and CDSs in the VSGnome. **(B)** PCA plot obtained from triplicated samples used for polyA-selected stranded RNA-seq. Sequence reads were mapped to CDSs in the Lister 427 genome.

**Supplementary Figure 5**

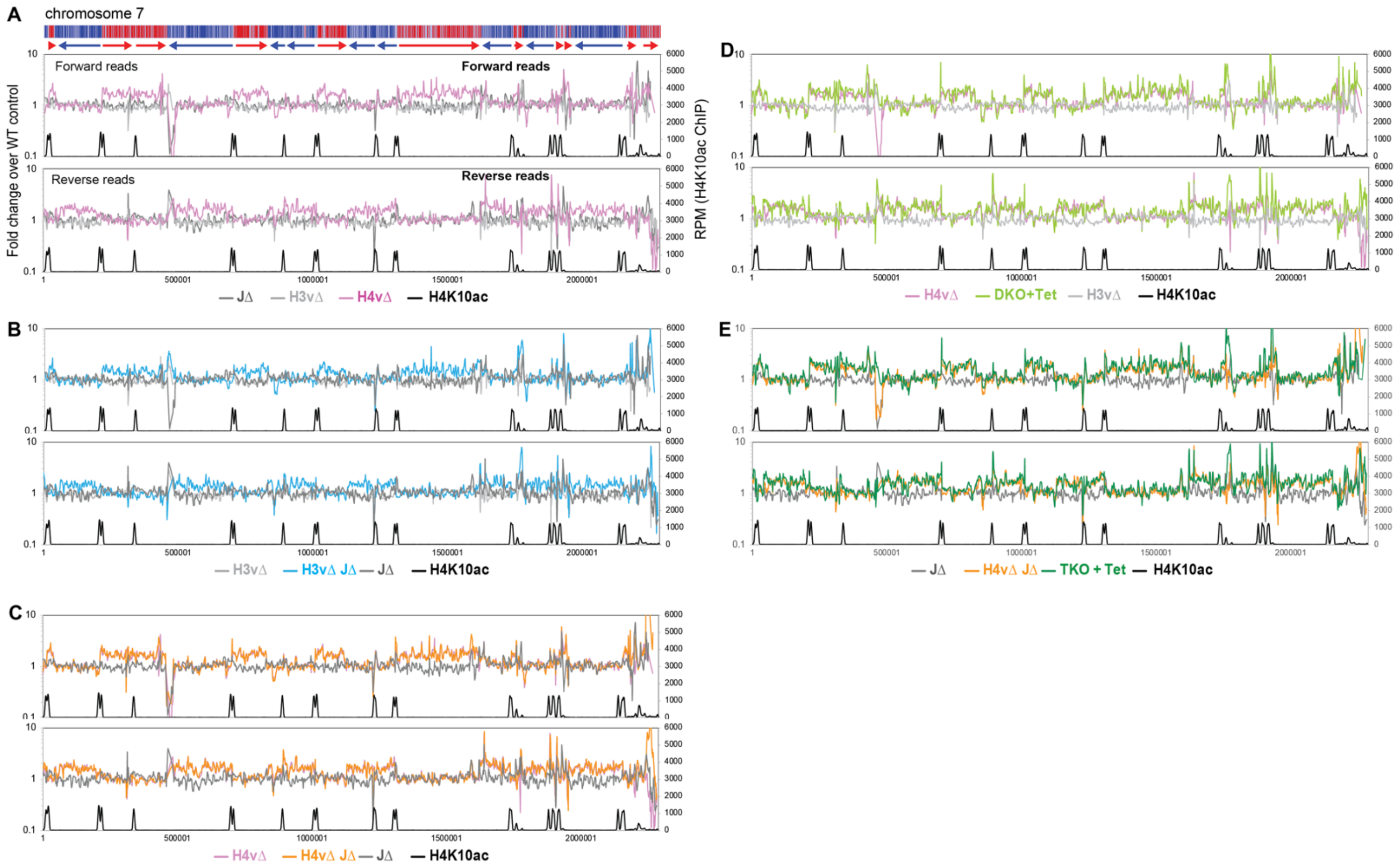


**Supplementary Figure 5. H4v is the major transcription termination signal and H3v-J has a secondary role**. Global transcription was examined by rRNA-depleted stranded RNA-seq and mapping the sequence reads to the *T. brucei* Lister 427 genome. Forward and reverse reads were analyzed separately with sliding windows (5kb bin, 1kb step) as in Figure 2. Chromosome 7 is shown as an example. Each plot compares level of transcripts between **(A)** JΔ, H4vΔ, and H3vΔ. **(B)** JΔ, H4vΔ, and H4vΔ JΔ. **(C)** JΔ, H3vΔ, and H3vΔ. **(D)** H3vΔ, H4vΔ, and Tet-treated DKO (H3vΔ H4vΔ). **(E)** JΔ, H4vΔ JΔ, Tet-treated TKO (H3vΔ H4vΔ JΔ).

Supplementary Figure 6

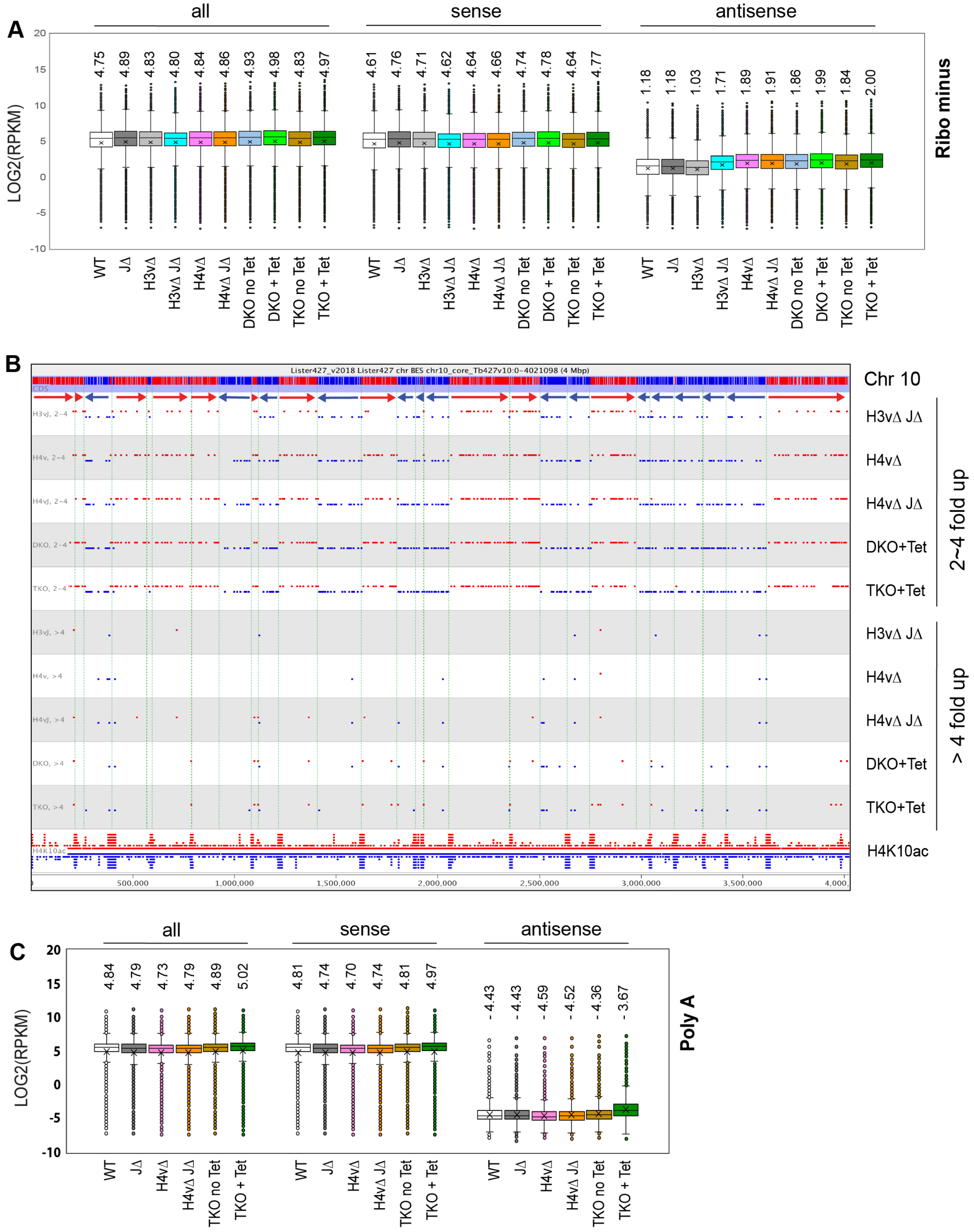


**Supplementary Figure 6. Increased antisense transcription levels in H4vΔ mutants.** $Log_2$(RPKM) values from all reads, reads mapping to opposite direction of genes (sense transcription) or reads mapping the same direction as genes (antisense transcription) were obtained for 8,428 CDSs. Box plots compare between wild type and KO mutant strains. **(A)** rRNA depleted stranded RNA-seq and **(C)** poly A selected stranded RNA seq samples. Mean values are shown and indicated as x. Outliers are shown as circles. Statistical analysis is summarized in Supplementary Table 7. **(B)** Location of CDSs in chromosome 10 that were 2~4-fold upregulated or more than 4-fold upregulated in H3vΔ JΔ, H4vΔ, H4vΔ JΔ, and Tet-treated DKO and TKO, compared to WT.

#### Supplementary Figure 7

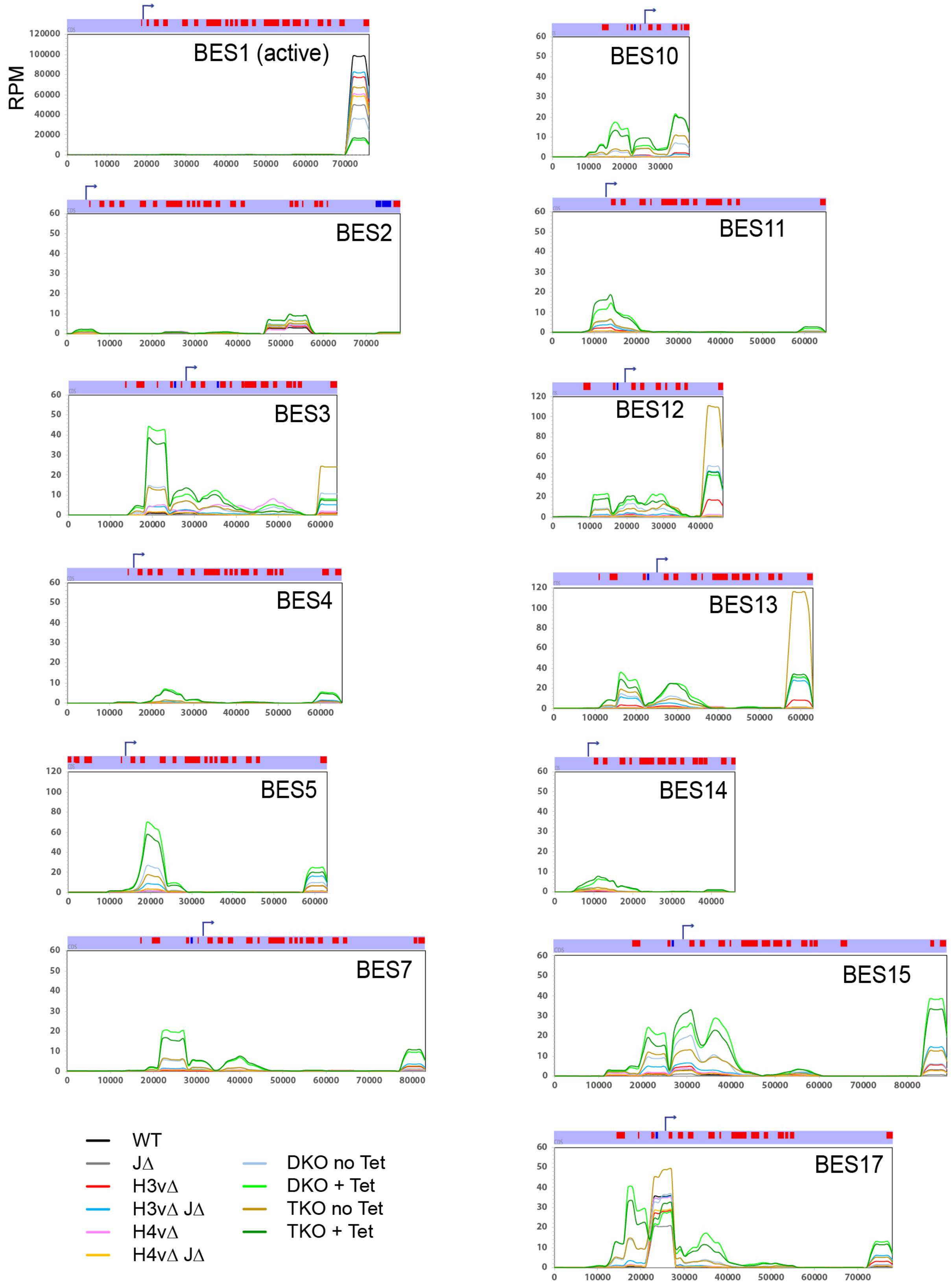


**Supplementary Figure 7. TTS chromatin marks are important in repression of promoter proximal and telomere proximal genes within BESs.** Lister 427 reference genome also contain BES sequence information. RPM values from reads mapping to BESs were plotted over each BES. Location of RNA pol I promoter is shown as arrows with bent tips.

### Supplementary Figure 8

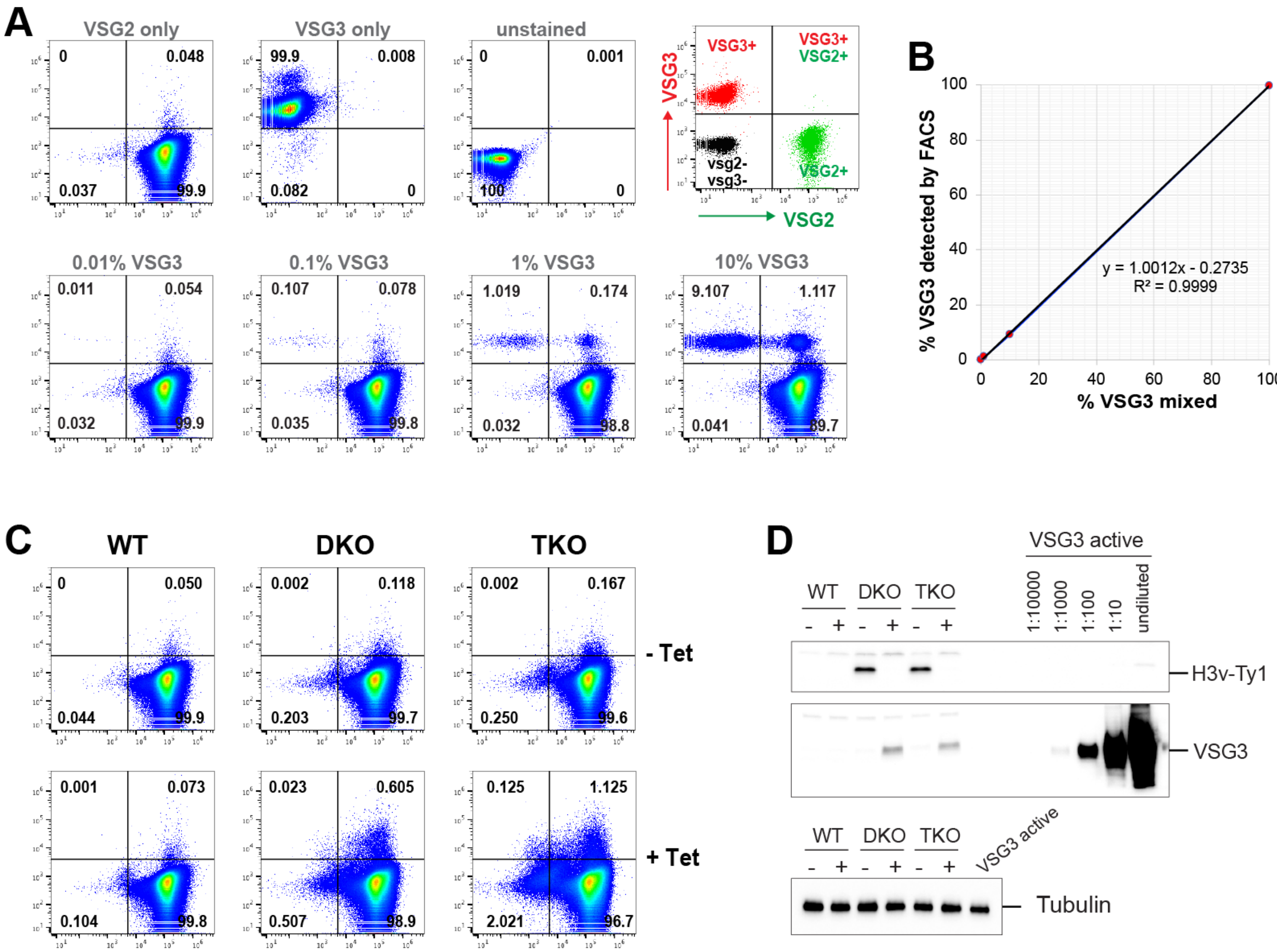


**Supplementary Figure 8. VSG switching in DKO and TKO strains. (A)** Detection of non-VSG2 expressing cells by flow cytometry. VSG2 expressing cells were mixed with VSG3 expressing cells at an indicated percentage. Live cells expressing VSG2 alone or VSG3 alone, or both were incubated with fluorophore conjugated VSG antibodies, anti-VSG2-Dylight 488 and anti-VSG2-Dylight 650, and then analyzed by flow cytometry. **(B)** Percent of VSG3 mixed in and percent of VSG3 detected by FACS. **(C)** WT, DKO and TKO cells treated with or without Tet were incubated with anti-VSG2-Dylight 488 and anti-VSG2-dylight 650, and then analyzed by flow cytometry. **(D)** Whole cells from (C) were collected and analyzed by western blot.

### Supplementary Figure 9

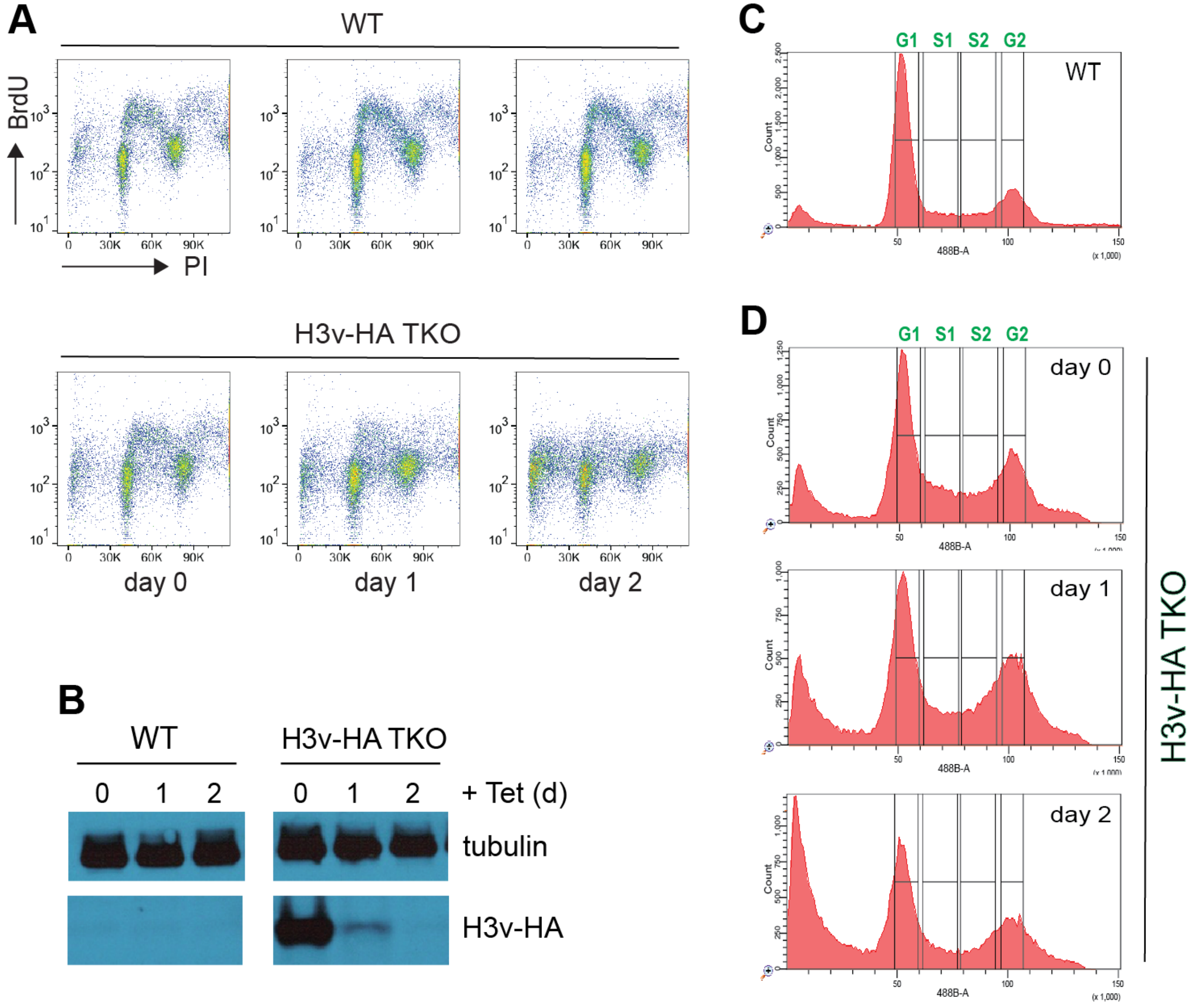


**Supplementary Figure 9. BrdU incorporation assay and cell-cycle sorting of wild type and triple KO mutants for MFA-seq experiment. (A)** BrdU incorporation assay. WT[32] and floxed H3v-HA TKO strain treated with tetracycline for 0, 1, and 2 days were pulse-labeled with 500 μM BrdU for 40 min. Bulk DNA was stained with PI, and cells were analyzed by flow cytometry. **(B)** Western blot confirming depletion of H3v-HA in the TKO strain after Tet treatment. Tubulin was used as a loading control. FACS sorting for MFA-seq assay; **(C)** WT; **(D)** TKO strain at day 0, 1, and 2 after tetracycline addition.

### Supplementary Figure 10

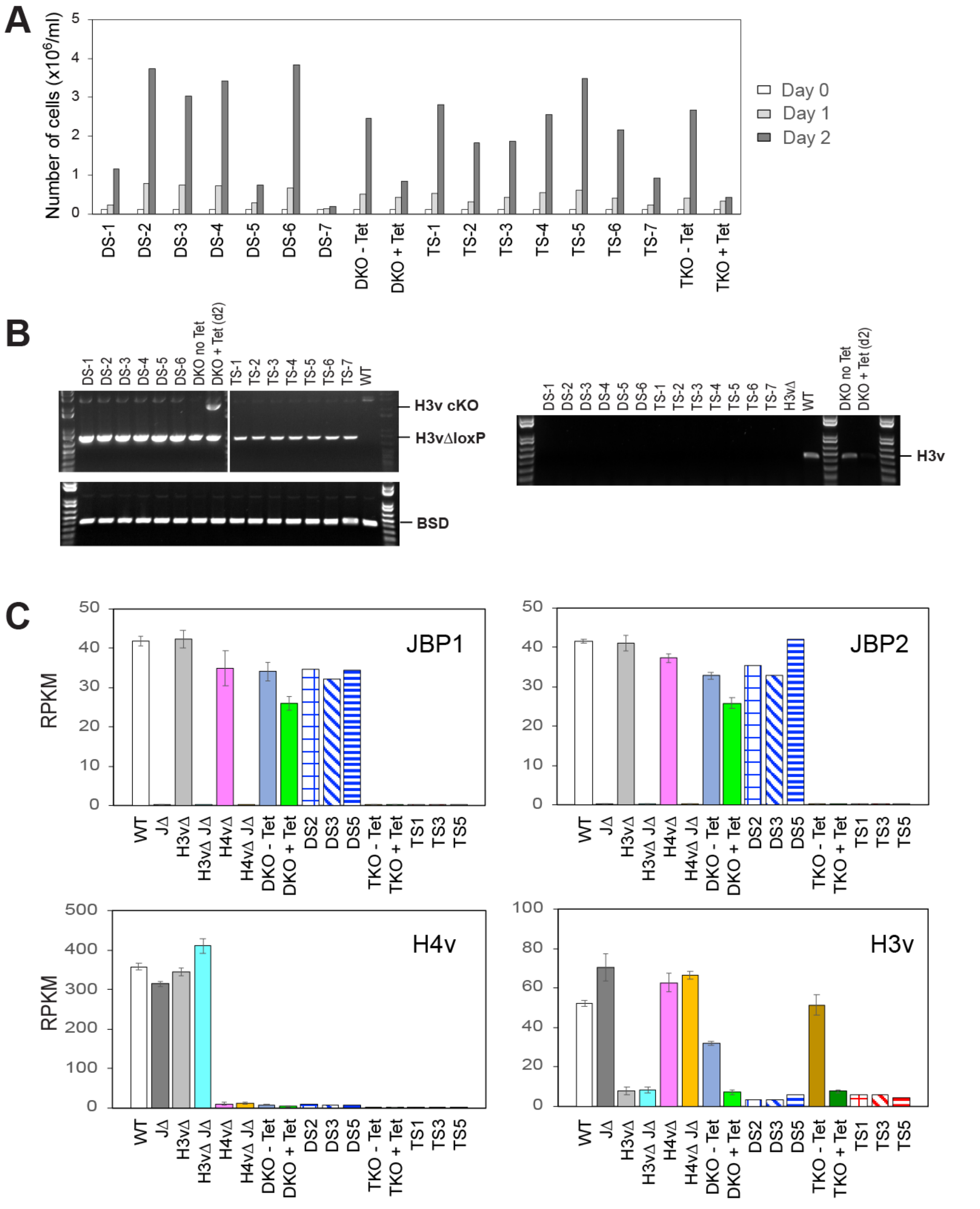


**Supplementary Figure 10. Control data for DS and TS clones. (A)** Growth of DS and TS clones. Surviving clones identified from the 96-well plate (7 DS and 7 TS clones) were monitored for cell growth. **(B)** PCR genotyping confirming the loss of H3v protein. **(C)** Levels of H3v, H4v, JBP1 and JBP2 RNA in the KO mutants, DS, and TS clones used in rRNA-depleted stranded RNA-seq. RPKM values of reads mapping to each gene in WT and mutants.

**Supplementary Figure 11**

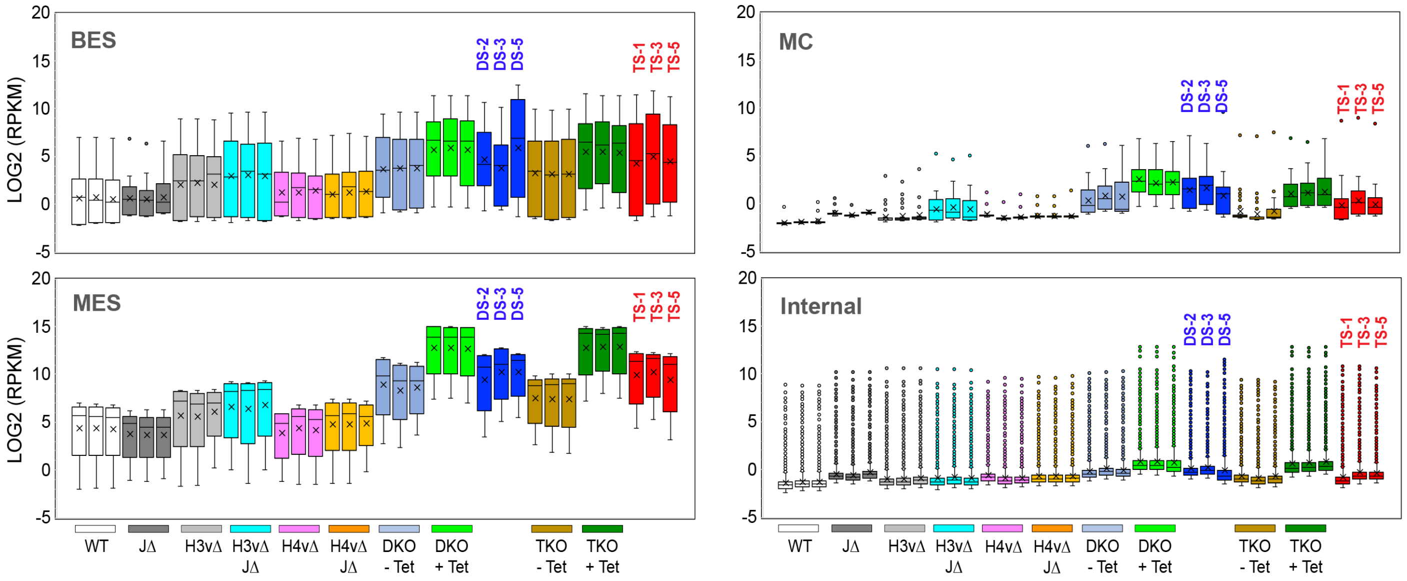


**Supplementary Figure 11.** Silencing of BES, MES, MC and chromosome-internal VSGs in DS and TS clones was compared with replicates of WT and KO mutant strains. Statistical analyses for box plots are summarized in Supplementary Table 9.

### Supplementary Figure 12

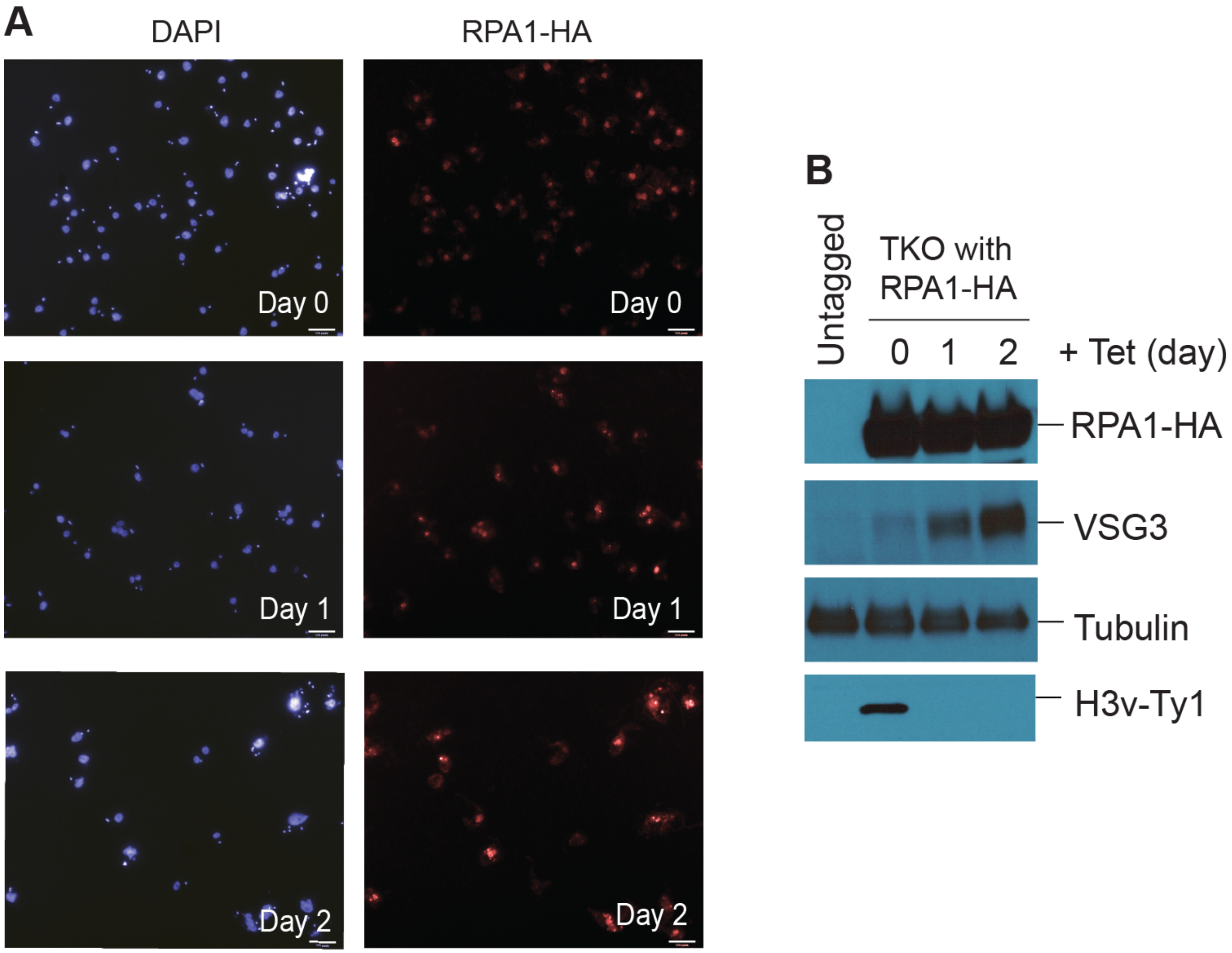


**Supplementary Figure 12. Formation of RPA1 nuclear foci in TKO cells. (A)** TKO strain expressing *Tb*RPA1 tagged with HA epitope was treated with Tet for 0, 1, and 2 days. Cells were then fixed and examined by immunofluorescence. Bulk DNA was stained with DAPI (blue) and *Tb*RPA1 with mouse anti-HA followed by secondary antibodies conjugated with Alexa 567(red). **(B)** Proteins were analyzed by western blot.

#### Supplementary Figure 13

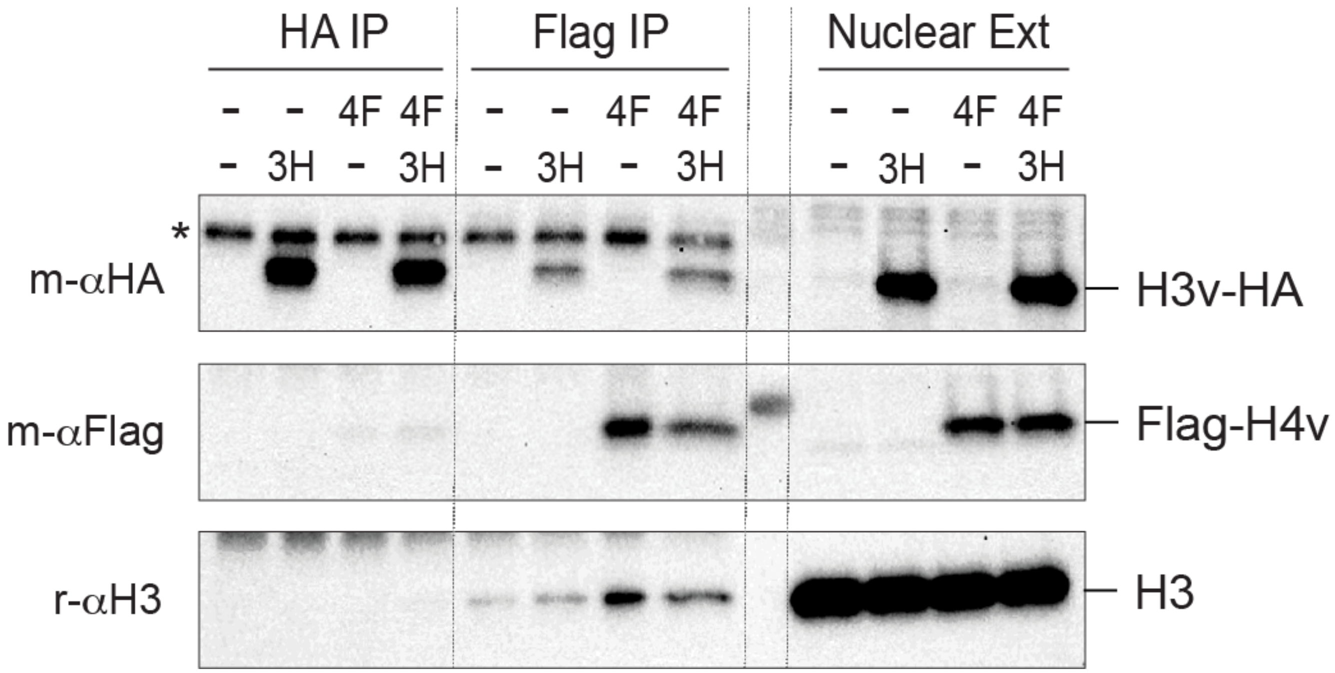


**Supplementary Figure 13. H3v and H4v are not in the same nucleosome.** Nuclear lysate was prepared from WT, a strain expressing H3v-HA ("3H") or Flag-H4v ("4F") only, or both. Lysates were immunoprecipitated with rabbit anti-HA or Flag antibodies and blots were probed with mouse anti-HA, Flag, or rabbit H3 antibodies. Asterisk is non-specific band.

#### Supplementary Tables

**Supplementary Table 1. List of cell lines, plasmids and oligo nucleotides used in this study**

A. List of cell lines used in this study
B. List of plasmids used in this study
C. List of oligo nucleotides used in this study

**Supplementary Table 2. Summary of Bowtie 2 mapping**

A. Stranded RNA-seq samples prepared with rRNA depletion and mapped to VSGnome
B. Stranded RNA-seq samples prepared with rRNA depletion and mapped to Lister 427
C. Stranded RNA-seq samples prepared with rRNA depletion and mapped to *Tb*927v5
D. Stranded RNA-seq samples prepared with polyA selection and mapped to Lister 427
E. MFA-seq reads mapped to Lister427

**Supplementary Table 3. Correlation between replicate samples from stranded RNA-seq**

A. rRNA-depleted stranded RNA-seq samples analyzed for Lister 427 CDSs
B. rRNA-depleted stranded RNA-seq samples analyzed for VSG CDSs
C. rRNA-depleted stranded RNA-seq samples analyzed for *Tb*927v5 Genes
D. PolyA-selected stranded RNA-seq samples analyzed for Lister 427 CDSs

**Supplementary Table 4. PCA analysis**

A. rRNA-depleted stranded RNA-seq samples analyzed for Lister 427 CDSs
B. rRNA-depleted stranded RNA-seq samples analyzed for VSG CDSs
C. rRNA-depleted stranded RNA-seq samples analyzed for *Tb*927v5 Genes
D. PolyA-selected stranded RNA-seq samples analyzed for Lister 427 CDSs

**Supplementary Table 5. rRNA-depleted stranded RNA-seq mapped to Lister 427 and analyzed with sliding window**

A. RPM values obtained from rRNA-depleted stranded RNA-seq samples analyzed with sliding window (5kb bin, 1kb step): Forward reads only
B. RPM values obtained from rRNA-depleted stranded RNA-seq samples analyzed with sliding window (5kb bin, 1kb step): Reverse reads only

**Supplementary Table 6. Transcriptome analyses**

A. $LOG_2$(RPKM) values of Lister 427 CDSs from rRNA-depleted stranded RNA-seq: all, sense and antisense reads (averaged)
B. $LOG_2$(RPKM) values of Lister 427 CDSs from rRNA-depleted stranded RNA-seq samples: all reads (all replicates)
C. $LOG_2$(RPKM) values of Lister 427 CDSs from rRNA-depleted stranded RNA-seq samples: sense reads only (all replicates)
D. $LOG_2$(RPKM) values of Lister 427 CDSs from rRNA-depleted stranded RNA-seq samples: antisense reads only (all replicates)
E. $LOG_2$(RPKM) values of Lister 427 CDSs from polyA-selected stranded RNA-seq samples: all, sense and antisense reads (averaged)
F. $LOG_2$(RPKM) values of *Tb*927 Genes from rRNA-depleted stranded RNA-seq samples: all reads (all replicates)
G. $LOG_2$(RPKM) values of *Tb*927 Genes from rRNA-depleted stranded RNA-seq samples: sense reads only (all replicates)
H. $LOG_2$(RPKM) values of *Tb*927 Genes from rRNA-depleted stranded RNA-seq samples: antisense reads only (all replicates)

**Supplementary Table 7. Statistics for transcriptome analyses for supplementary Table 6**

A. Lister 427 CDSs from rRNA-depleted stranded RNA-seq samples
B. Lister 427 CDSs from polyA-selected stranded RNA-seq samples
C. *Tb*927v5 Genes from rRNA-depleted stranded RNA-seq samples

**Supplementary Table 8. PolyA-selected stranded RNA-seq mapped to Lister 427 and analyzed with sliding window**

A. RPM values obtained from polyA-selected stranded RNA-seq samples analyzed with sliding window (5kb bin, 1kb step): Forward reads only
B. RPM values obtained from polyA-selected stranded RNA-seq samples analyzed with sliding window (5kb bin, 1kb step): Reverse reads only

**Supplementary Table 9. VSGnome mapping, analysis and statistics**

A. $LOG_2$(RPKM) value of all VSGs in all replicates of WT and 9 KO mutant strains and clones of DS and TS
B. Statistical analysis for MES VSG expression (all replicates)
C. Statistical analysis for BES VSG expression (all replicates)
D. Statistical analysis for MC VSG expression (all replicates)
E. Statistical analysis for Chromosome internal VSG expression (all replicates)
F. $LOG_2$(RPKM) of all VSGs (averaged)
G. Heat map comparison between WT and KO mutants and between clones of DS & TS and their parental strains.
H. Statistical analysis for MC VSG expression (averaged)
I. Statistical analysis for Chromosome Internal VSGs (averaged)

**Supplementary Table 10. MFA-seq mapped to Lister 427 and analyzed with sliding window (10kb bin, 2.5kb step, RPM value)**

**Supplementary Table 11. rRNA-depleted stranded RNA-seq mapped to Lister 427 and analyzed with sliding window for DS and TS clones**

A. RPM values obtained from rRNA-depleted stranded RNA-seq samples analyzed with sliding window (10kb bin, 2.5kb step): Forward reads only
B. RPM values obtained from rRNA-depleted stranded RNA-seq samples analyzed with sliding window (10kb bin, 2.5kb step): Reverse reads only
